# Neutralizing and protective human monoclonal antibodies recognizing the N-terminal domain of the SARS-CoV-2 spike protein

**DOI:** 10.1101/2021.01.19.427324

**Authors:** Naveenchandra Suryadevara, Swathi Shrihari, Pavlo Gilchuk, Laura A. VanBlargan, Elad Binshtein, Seth J. Zost, Rachel S. Nargi, Rachel E. Sutton, Emma S. Winkler, Elaine C. Chen, Mallorie E. Fouch, Edgar Davidson, Benjamin J. Doranz, Robert H. Carnahan, Larissa B. Thackray, Michael S. Diamond, James E. Crowe

## Abstract

Most human monoclonal antibodies (mAbs) neutralizing SARS-CoV-2 recognize the spike (S) protein receptor-binding domain and block virus interactions with the cellular receptor angiotensin-converting enzyme 2. We describe a panel of human mAbs binding to diverse epitopes on the N-terminal domain (NTD) of S protein from SARS-CoV-2 convalescent donors and found a minority of these possessed neutralizing activity. Two mAbs (COV2-2676 and COV2-2489) inhibited infection of authentic SARS-CoV-2 and recombinant VSV/SARS-CoV-2 viruses. We mapped their binding epitopes by alanine-scanning mutagenesis and selection of functional SARS-CoV-2 S neutralization escape variants. Mechanistic studies showed that these antibodies neutralize in part by inhibiting a post-attachment step in the infection cycle. COV2-2676 and COV2-2489 offered protection either as prophylaxis or therapy, and Fc effector functions were required for optimal protection. Thus, natural infection induces a subset of potent NTD-specific mAbs that leverage neutralizing and Fc-mediated activities to protect against SARS-CoV-2 infection using multiple functional attributes.

## INTRODUCTION

Since the emergence of severe acute respiratory syndrome coronavirus 2 (SARS-CoV-2) as a major threat to global public health, many studies have focused efforts on discovery of potent neutralizing monoclonal antibodies (mAbs) against the spike (S) protein of SARS-CoV-2 (Baum et al., 2020a; Cao et al., 2020; Hansen et al., 2020; Ju et al., 2020; Liu et al., 2020a; Pinto et al., 2020; Robbiani et al., 2020; Zost et al., 2020b). The S protein exists as a trimer on the surface of SARS-CoV-2 and facilitates entry of the virus into cells. The receptor binding domain (RBD) situated in the S1 region of the S protein binds to human angiotensin-converting enzyme 2 (hACE2). The S2 region then changes conformation and inserts its fusion peptide into the target cell membrane, thus triggering viral fusion and entry. Previous studies have demonstrated that the RBD region is a key target for potently neutralizing antibodies (Alsoussi et al., 2020; Barnes et al., 2020; Baum et al., 2020a; Hansen et al., 2020; Hassan et al., 2020; Zost et al., 2020a). These studies also defined inhibition of S trimer binding to the cellular receptor ACE2 as a principal mechanism of action of RBD-targeting antibodies against SARS-CoV-2, and showed protection in animals against infection and disease by this class of mAbs (Baum et al., 2020a; Hansen et al., 2020; Hassan et al., 2020; Zost et al., 2020a).

Since the start of the outbreak, circulating SARS-CoV-2 field strains have acquired genetic changes that facilitate transmission (Avanzato et al., 2020; Choi et al., 2020; Hou et al., 2020; Kemp et al., 2020; McCarthy et al., 2020; Plante et al., 2020; Rambaut et al., 2020; Tegally et al., 2020). This rapid viral evolution also could affect the protective efficacy of vaccines and mAb-based therapeutics currently in clinical trials or approved under emergency use authorization. Most of the mAbs under evaluation in clinical trials or authorized for the emergency use bind to the RBD (Baum et al., 2020a; Zost et al., 2020a). Several groups have described mAbs targeting non-RBD regions, but their epitopes, mechanisms of action, and protective activity *in vivo* remain unclear (Chi et al., 2020; Liu et al., 2020a). Here we define the structure-function relationship of potent NTD-reactive antibodies from a panel of 389 human SARS-CoV-2 S protein mAbs we isolated from survivors of natural infection (Zost et al., 2020a; Zost et al., 2020b). We found 43 mAbs recognizing the NTD. Three of the 43 NTD-reactive mAbs exhibited neutralizing capacity against authentic SARS-CoV-2 virus (Zost et al., 2020b), with two being potently inhibitory. We mapped the epitopes for the two most potently neutralizing NTD-reactive mAbs and dissected the mechanism by which these mAbs inhibited SARS-CoV-2 infection. These two mAbs conferred protection in hACE2-expressing mice when administered either as prophylaxis or therapy, and intact Fc effector functions were required for optimal activity *in vivo*. Thus, we show that regions of the NTD are recognized by neutralizing and protective antibodies against SARS-CoV-2 and could function as part of antibody cocktails to minimize the selection of escape variants or resistance to natural variants in RBD as they emerge.

## RESULTS

### Two non-RBD anti-SARS-CoV-2 S-protein-specific mAbs potently neutralize virus

We previously isolated 389 SARS-CoV-2 reactive mAbs from the B cells of convalescent COVID-19 patients. While most of the neutralizing antibodies mapped to the RBD and blocked ACE2 binding to the S protein, multiple neutralizing mAbs that did not bind RBD were also identified (Zost et al., 2020a; Zost et al., 2020b). We selected the two most potently neutralizing S-protein-reactive mAbs that did not bind to RBD, designated COV2-2676 and COV2-2489, for further study. We first assessed binding and neutralizing activities of COV2-2676 and COV2-2489 to determine their potency. Both mAbs bound weakly to a recombinant SARS-CoV-2 S6_Pecto_ protein expressed as a trimer, but failed to react with a recombinant soluble NTD protein. In contrast, additional SARS-CoV-2-specific non-RBD mAbs, COV2-2263 and COV2-2490, which are non-neutralizing(Zost et al., 2020a), reacted to both SARS-CoV-2 S6_Pecto_ protein and recombinant soluble NTD protein, whereas the dengue virus mAb DENV-2D22, a negative control for the assays, reacted to neither (**Fig 1A** and **B**). We performed two types of virus neutralization assays with COV2-2676 and COV2-2489. The first method used authentic WA1/2020 strain SARS-CoV-2 in a focus reduction neutralization test (FRNT) (Harcourt et al., 2020a; Harcourt et al., 2020b), whereas the second was a real-time cell analysis (RTCA) assay using recombinant replication-competent vesicular stomatitis virus (VSV) expressing the SARS-CoV-2 S protein in place of the endogenous VSV glycoprotein (Case et al., 2020b). Both COV2-2676 and COV2-2489 individually neutralized the authentic SARS-CoV-2 in a dose-dependent manner with a half-maximal inhibitory (IC_50_) value of 501 or 199 ng/mL, respectively (**Fig 1C**). In addition, COV2-2676 and COV2-2489 neutralized chimeric VSV-SARS-CoV-2 with IC_50_ values of 38 or 56 ng/mL, respectively (**Fig 1D**).

**Figure 1.**
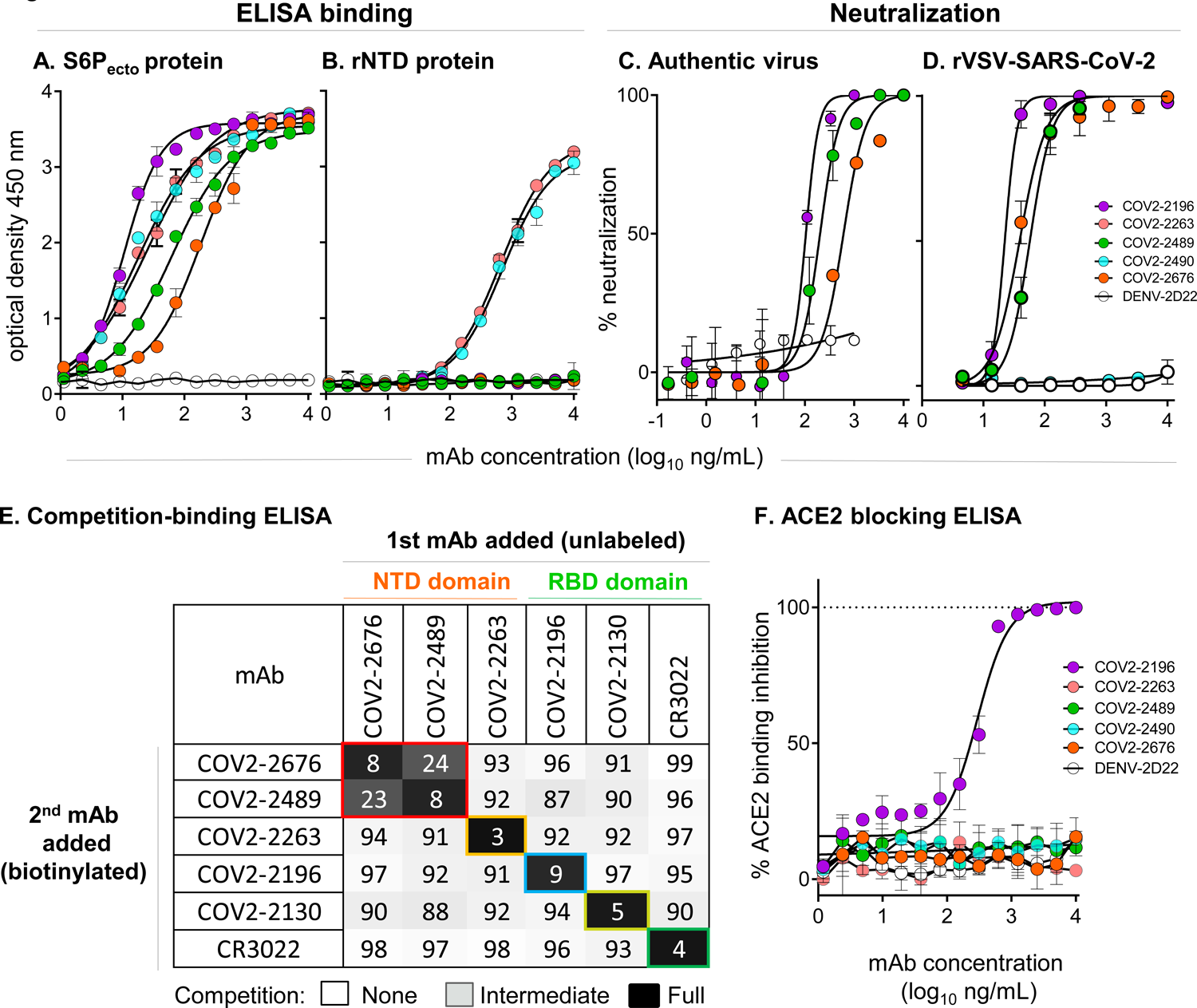
Two non-RBD-reactive mAbs specific to SARS-CoV-2 S protein neutralize the virus. **A.** ELISA binding of COV2-2196, COV2-2263, COV2-2489, COV2-2490, COV2-2676 or rDENV-2D22 to trimeric S-6_Pecto_. Data are mean ± standard deviatioms (S.D.) of technical triplicates from a representative experiment repeated twice. **B.** ELISA binding of COV2-2196, COV2-2263, COV2-2489, COV2-2490, COV2-2676 or rDENV-2D22 to rNTD protein. Data are mean ± S.D. of technical triplicates from a representative experiment repeated twice. B. Neutralization curves for COV2-2196, COV2-2489, COV2-2676 or rDENV-2D22 using wild-type SARS-CoV-2 in a FRNT assay. Error bars indicate S.D.; data represent at least two independent experiments performed in technical duplicate. C. Neutralization curves for COV2-2196, COV2-2489, COV2-2490, COV2-2676 or rDENV-2D22 in a SARS-CoV-2-rVSV neutralization assay using RTCA. Error bars indicate S.D.; data are representative of at least two independent experiments performed in technical duplicate. D. Competition binding of the panel of neutralizing mAbs with reference mAbs COV2-2130, COV2-2196, COV2-2263, COV2-2489, COV2-2676 or rCR3022. Binding of reference mAbs to trimeric S-6_Pecto_ protein was measured in the presence of saturating concentration of competitor mAb in a competition ELISA and normalized to binding in the presence of rDENV-2D22. Black indicates full competition (<25% binding of reference antibody); grey indicates partial competition (25 to 60% binding of reference antibody); white indicates no competition (>60% binding of reference antibody). E. Human-ACE2-blocking curves for COV2-2196, COV2-2263, COV2-2489, COV2-2490, COV2-2676 and rDENV-2D22 in a human-ACE2-blocking ELISA. Data are mean ± S.D. of technical triplicates from a representative experiment repeated twice.

Competition-binding analysis using the S6_Pecto_ protein revealed that COV2-2676 and COV2-2489 competed for binding with one another (**Fig S1**). However, these mAbs did not compete for binding with any of the other 33 previously identified non-neutralizing NTD-reactive human mAbs (Zost et al., 2020b) or with mAbs that recognize non-overlapping antigenic sites on the surface of RBD (including COV2-2196, COV2-2130, and a recombinant form of CR3022(ter Meulen et al., 2006) [**Fig 1E**]). These findings indicate an antigenic site for COV2-2676 and COV2-2489 distinct from those previously identified for neutralizing human mAbs targeting the RBD. Moreover, neither COV2-2676 nor COV2-2489 inhibited the interaction of soluble ACE2 with soluble RBD protein (**Fig 1F**).

### COV2-2676 and COV2-2489 recognize the NTD of SARS-CoV-2 S protein

To determine the binding sites for these mAbs, we used negative-stain electron microscopy to image a stabilized trimeric form of the ectodomain of S protein (S6_Pecto_ trimer) in complex with Fab fragment forms of COV2-2676 or COV2-2489 (**Fig 2A**). These antibodies bind to the NTD and recognize the ‘closed’ conformational state of the S6_Pecto_ trimer. By overlaying the negative stain EM maps of the two Fab/S complexes, we determined that these antibodies bind to a common antigenic site on the NTD of the S6_Pecto_ trimer. These findings are consistent with the competition-binding data between COV2-2676 and COV2-2489. Sequence alignment of the variable gene sequences of these mAbs with previously published anti-NTD neutralizing mAbs 4A8 and 4-8 showed that COV2-2676 or COV2-2489 are independent clonotypes. COV2-2489 is encoded by heavy chain variable gene segment *IGHV-4-39.* COV2-2676 mAb is encoded by *IGHV1-69*, as is mAb 4-8, but the HCDR3 regions of the two mAbs differ completely, and mAb 4A8 is encoded by *IGHV1-24* (**Table S1**). We superimposed the COV2-2676 negative stain-EM Fab complex with the cryo-EM structure of mAb 4-8 and found that the binding interfaces of both mAbs are similar.

**Figure 2.**
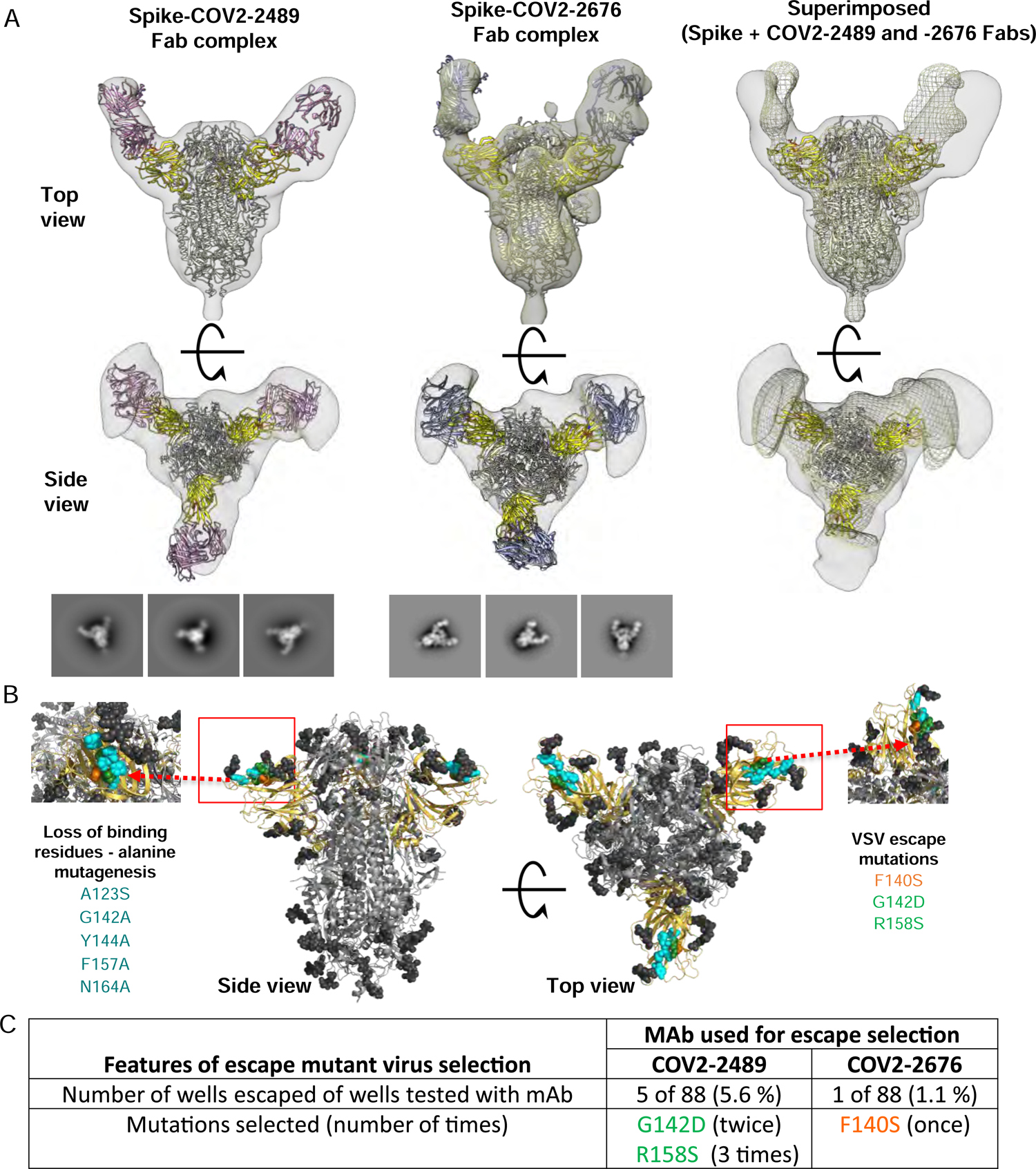
COV2-2676 and COV2-2489 bindng map to the NTD of SARS-COV-2-S protein. **A.** Top row (side view), bottom row (top view) of Fab–S6_Pecto_ closed trimer (S protein model PDB:7JJI) complexes visualized by negative-stain electron microscopy for COV2-2676 Fab model in pink, COV2-2489 Fab model in blue and superimpose 3D volume of CoV2-S-Fab 2676 complex in grey and CoV2-S-Fab 2489 in mesh. The S-NTD is shown in yellow and electron density in grey. Representative two-dimensional (2D) class averages for each complex are shown at the bottom (box size is 128 pixels, with 4.36 Å per pixel). Data are from a single experiment; detailed collection statistics are provided in **Supplementary Table 2**. **B.** Identification of critical contact residues by alanine-scanning mutagenesis. Top (side view) with loss of binding residues (cyan) for COV2-2489 or COV2-2676 to mutant S-NTD constructs, normalized to the wild-type. Bottom, escape mutations mapped to the NTD region for COV2-2489 (green G142D, R158S) or COV2-2676 (orange F140S). **C**. Results of viral selections with COV2-2489 or COV2-2676 individual mAbs. The number of replicates in which escape variants were selected is indicated. Mutations present in the NTD of the selected escape variants are indicated.

The heavy chain of the antibodies interact with the N3 and N5 loops of NTD (**Fig S2**). This revealed a distinct site of vulnerability on NTD region of spike protein for human neutralizing mAbs and suggested a convergent responses in SARS-CoV-2 immune individuals.

We next defined the antibody epitopes at the amino acid level using two complementary methods: alanine-scanning loss-of-binding experiments in cell-surface antigen display and selection of virus escape mutants followed by sequence analysis. Screening of the NTD Ala-scan library identified residues A123, G142, Y144, F157 and N164 as important for binding of COV2-2489, and Y144 for binding of mAb COV2-2676. None of these single-residue alanine mutants affected binding of the control NTD-reactive mAb COV2-2305, likely due to the location of key contact residues in the N3 and N5 loops of NTD (**Fig 2B** and **S3A**).

To identify neutralization escape mutations, we used a high-throughput RTCA assay as previously described (Gilchuk et al., 2020a; Gilchuk et al., 2020b). We selected viral variants that escape antibody neutralization at a single saturating concentration of 5 μg/mL (COV2-2676) or 50 μg/mL (COV2-2489) and identified point mutations. The mutations were F140S (orange) for COV2-2676, and G142D and R158S (green) for COV2-2489 (**Fig 2C**). We confirmed that the mutations in the viral variants we selected in the RTCA assay experiments above conferred resistance to 10 or 100 μg/mL of the rescpetive mAbs (**Fig S3B**). The location of the escape mutations was in the same region identified by the alanine-scanning method, although the specific amino acids differed between the two methods. Thus, the two neutralizing NTD antibodies bind to a common antigenic site, but the fine specificity of the epitopes differ.

### COV2-2676 and COV2-2489 bind avidly to S protein on the surface of infected cells, show cell-type specific neutralization, and inhibit at a post-attachment step

We evaluated the ability of mAbs targeting the NTD to bind to the surface of SARS-CoV-2-infected cells since this property could contribute to immune cell-mediated clearance *in vivo*. Following SARS-CoV-2 infection, cells were incubated with serial dilutions of COV2-2489, COV2-2676, the neutralizing RBD-targeting mAb COV2-2381(Zost et al., 2020a), or the dengue virus mAb 2D22 isotype control, prior to analysis of staining intensity by flow cytometry. COV2-2489 and COV2-2676 exhibited similar avidity for cell-surface-associated S protein, with EC50 values for staining of infected cells of 896 ng/mL or 1,438 ng/mL, respectively (**Fig 3A**, left panel). Whereas the RBD-targeting mAb COV2-2381 had a more potent EC50 value for binding to infected cells (45 ng/mL), the NTD-targeting mAbs had greater staining intensity of infected cells, as the peak integrated mean fluorescence intensity (iMFI) was higher for COV2-2489 and COV2-2676 than for COV2-2381 (**Fig 3A**, left and right panels, and **Fig S4A**). These data suggest that NTD-targeting mAbs can bind efficiently and at high density to S protein on the surface of SARS-CoV-2 infected cells. We next assessed the neutralizing activity of COV2-2489 and COV2-2676 by FRNT on additional cell lines: MA104 cells, HEK-293T cells ectopically expressing ACE2 (293T+ACE2), and Vero cells ectopically expressing TMPRSS2 (Vero+TMPRSS2), in comparison to wild-type Vero cells. For both mAbs, neutralization potency was greater on Vero cells than MA104 cells or 293T+ACE2 cells; both mAbs had weak neutralization potency when assessed using the latter two cell types (**Fig 3B**). TMPRSS2 over-expression did not alter the potency of the mAbs, as neutralization assays using Vero+TMPRSS2 cells had similar potency to the parental wild-type cell (**Fig 3B**). These data indicate that mAbs against the NTD exhibit cell type-dependent neutralization, which may be due to variable expression of entry factors and receptors on the surface of different cell types.

**Figure 3.**
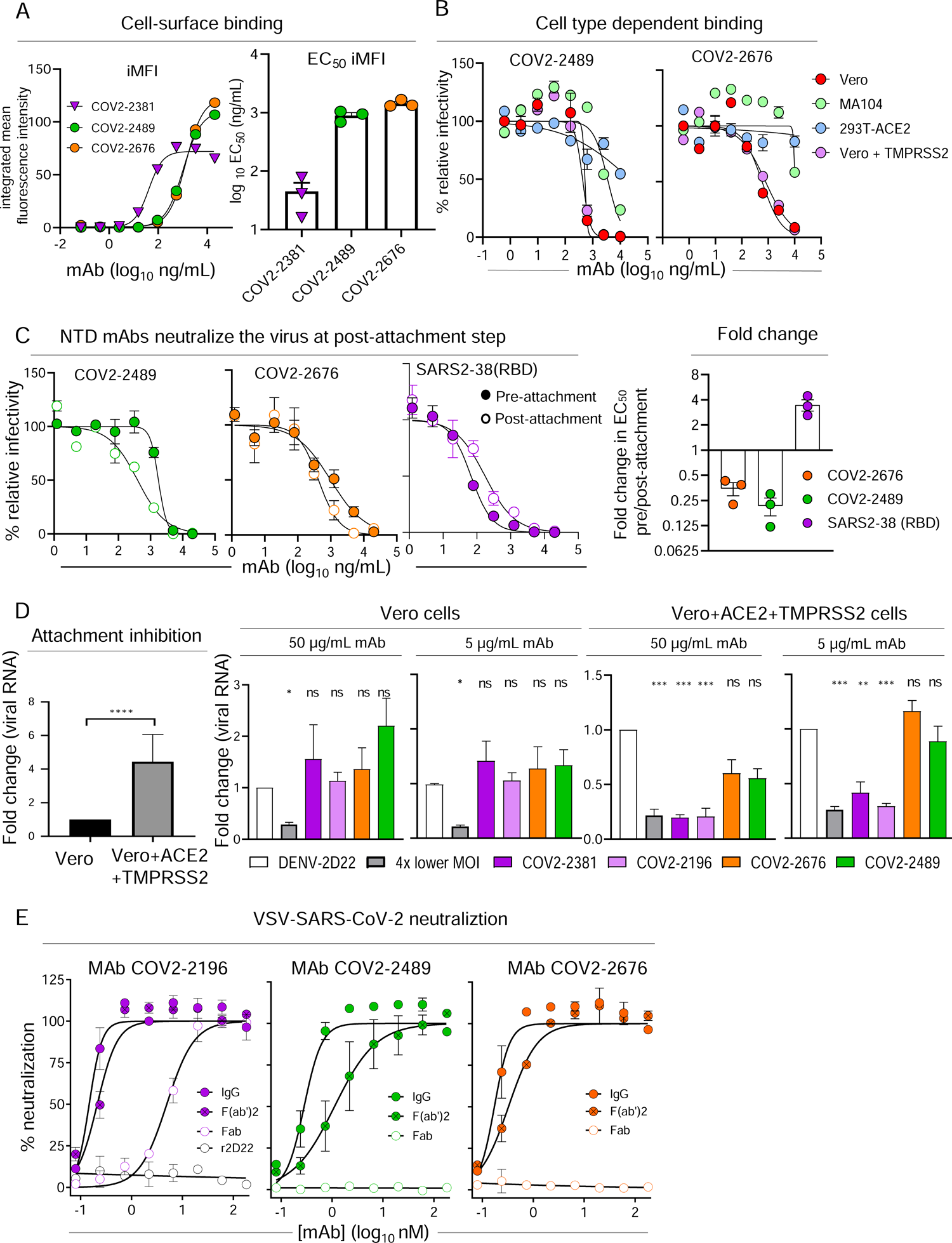
Virus neutralization and binding of infected cells by anti-SARS-CoV-2 mAbs targeting the NTD. **A.** SARS-CoV-2-infected Vero cells were stained with serial dilutions of COV2-2489, COV2-2676, COV2-2381, or DENV2-2D22 (isotype control) prior to analysis of staining intensity by flow cytometry. The positively stained cells were gated using uninfected and isotype control mAb-stained infected cells, and the integrated mean fluorescence intensity (iMFI) was determined by MFI of positive cells multiplied by the percent of total positive cells. The left panel shows representative dose response curves for the staining intensity of infected cells by each mAb. The right panel shows the mean EC50 values for infected cell staining, determined from three independent experiments. Error bars represent SEM. **B-D.** The neutralization potency of COV2-2489 and COV2-2676 against SARS-CoV-2 was assessed by FRNT using (**B**) Vero, MA104, 293T+ACE2, and Vero+TMPRSS2 cells. Results are representative of three independent experiments performed in duplicate. (**C**) COV2-2489 and COV2-2676 were assayed for neutralization potency by modified FRNT in which mAb was added to SARS-CoV-2 before (pre-attachment, filled cirlces) or after (post-attachment, open circles) virus was absorbed to Vero E6 cells. (**C**) Error bars represent the range from two technical replicates. Data shown are representative of three independent experiments. **D**. Attachment inhibition. (*Left panel*) Vero or Vero+hACE2+TMPRSS2 cells were incubated with SARS-CoV-2 at 4°C for 1 h. Afer extensive washing, cell-bound viral RNA was measured by qRT-PCR. (*Right 4 panels*) SARS-CoV-2 was pre-incubated with 5 or 50 μg/mL of indicated anti-RBD or anti-NTD mAbs for 1 h prior to addition to Vero or Vero+hACE2+TMPRSS2 cells. Cell-bound viral RNA was measured by qRT-PCR. Data are pooled from three independent experiments. (*Left*) t-test: ****p<0.0001; (*Right*) One-way ANOVA with Dunnett’s multiple comparisons test compared to isotype control mAb treatment: *p<0.05; **p<0.01; ***p<0.001. **E**. Neutralization curves for COV2-2196 IgG, F(ab′)_2_, F(ab) or rDENV-2D22; COV2-2489 IgG, F(ab′)_2_, F(ab); or COV2-2676 IgG, F(ab′)_2_, F(ab) in a SARS-CoV2-rVSV assay using RTCA. Error bars indicate S.D.; data represent at least two independent experiments performed in technical triplicates.

To probe further the mechanism of neutralization of mAbs targeting the NTD, we performed modified FRNTs, in which SARS2-CoV-2 was incubated with mAb before (pre-attachment FRNT) or after (post-attachment FRNT) virus was absorbed to the surface of Vero cells. Both COV2-2489 and COV2-2676 efficiently neutralized SARS-CoV-2 when added following virus absorption to the surface of cells (**Fig 3C**). These data suggest that mAbs targeting the NTD can neutralize SARS-CoV-2 at a post-attachment step of the viral life cycle in Vero cells.

We also performed experiments that directly assessed the ability of NTD mAbs to block virus attachment to Vero E6 cells and Vero overexpressing hACE2 and TMPRSS2 cells. When hACE2 and TMPRSS2 were expressed in cells ectopically, greater SARS-CoV-2 attachment was observed (**Fig 3D**, left panel). Neither anti-NTD (COV2-2489 or COV2-2676) nor anti-RBD neutralizing mAbs (COV2-2381 or COV2-2196) blocked attachment to Vero E6 cells (**Fig 3D**, middle panels). However, in hACE2 and TMPRSS2 over expressing cells binding was inhibited by anti-RBD but not anti-NTD mAbs (**Fig 3D**, right panels).

### COV2-2676 and COV2-2489 require intact IgG or F(ab′)_2_ for neutralizing activity

Next, to determine the valency of binding required for activity, we created recombinant IgG, F(ab′)_2_, and Fab forms of COV2-2676 and COV2-2489 and investigated their neutralizing potency on Vero cells. IgG or F(ab′)_2_ versions of COV2-2676 or COV2-2489 exhibited neutralizing activity in the RTCA assay, with IC_50_ values of 0.17 or 0.32 nM (COV2-2676) or 0.28 or 1.20 nM (COV2-2489), respectively (**Fig 3E**). Neutralizing activity of the F(ab′)_2_ molecules was comparable to that of the IgG versions. However, Fab fragments of COV2-2676 or COV2-2489 did not neutralize infection, whereas a potently neutralizing RBD antibody did inhibit infection as a Fab (**Fig 3E**). The reason for loss of neutralization activity by Fab fragments of COV2-2676 and COV2-2489 is unclear. The larger IgG or F(ab′)_2_ might sterically hinder functional interactions of the S protein in entry. Alternatively, the monovalent Fab molecule may bind with too low an affinity to the NTD to inhibit entry.

### COV2-2676 and COV2-2489 protect against SARS-CoV-2 challenge in mice

We tested the protective efficacy of COV2-2676 or COV2-2489 monotherapy in a K18-hACE2-transgenic mouse model of SARS-CoV-2 infection (Golden et al., 2020; Oladunni et al., 2020; Winkler et al., 2020). Mice treated with 200 µg (10 mg/kg) of the isotype control mAb DENV-2D22 one day before intranasal inoculation with 10^3^ plaque-forming units (PFU) of SARS-CoV-2 (strain 2019n-CoV/USA_WA1/2020) experienced weight loss between 4 and 7 days after inoculation (**Fig 4A**). In contrast, prophylaxis with 200 µg of COV2-2676 or COV2-2489 prevented weight loss in all mice, in a manner similar to that mediated by COV2-2381, a neutralizing RBD-specific human mAb known to protect *in vivo* against SARS-CoV-2 infection (Zost et al., 2020a). Pre-treatment with COV2-2676 or COV2-2489 also significantly decreased viral burden at 7 days post-infection (dpi) in the upper and lower respiratory tracts and at a distal site, the heart, compared to DENV-2D22, in a manner similar to COV2-2381 (**Fig 4B**). Prophylaxis with 200 µg of COV2-2676, COV2-2489, or COV2-2381 was associated with serum IgG1 concentrations of ∼ 4 µg/mL (**Fig 4C**). Pre-treatment with COV2-2676 or COV2-2489 protected against weight loss and viral burden at a 25-fold lower dose, whereas pre-treatment with COV2-2381 protected at a 125-fold lower dose. These treatments were associated with serum IgG1 concentrations of ∼200 ng/mL or ∼10 ng/mL, respectively (**Fig 4D-F**). Since SARS-CoV-2 infection causes substantial lung inflammation in K18-hACE2 mice and in humans (Golden et al., 2020; Oladunni et al., 2020; Winkler et al., 2020), we evaluated the effects of mAb treatment on cytokine and chemokine production in lung tissues at 7 dpi. Prophylaxis with 200 µg of COV2-2676 reduced multiple cytokine and chemokine levels in a manner similar to COV2-2381, in contrast with results from the isotype control mAb DENV-2D22 group (**Fig 4G** and **S5A**). Consistent with these results, analysis of hematoxylin and eosin-stained lung sections showed a reduction in perivascular and parenchymal immune cell infiltration, and alveolar space consolidation in the lungs of COV2-2676 or COV2-2381-treated mice compared to DENV-2D22-treated animals (**Fig 4H**).

**Figure 4.**
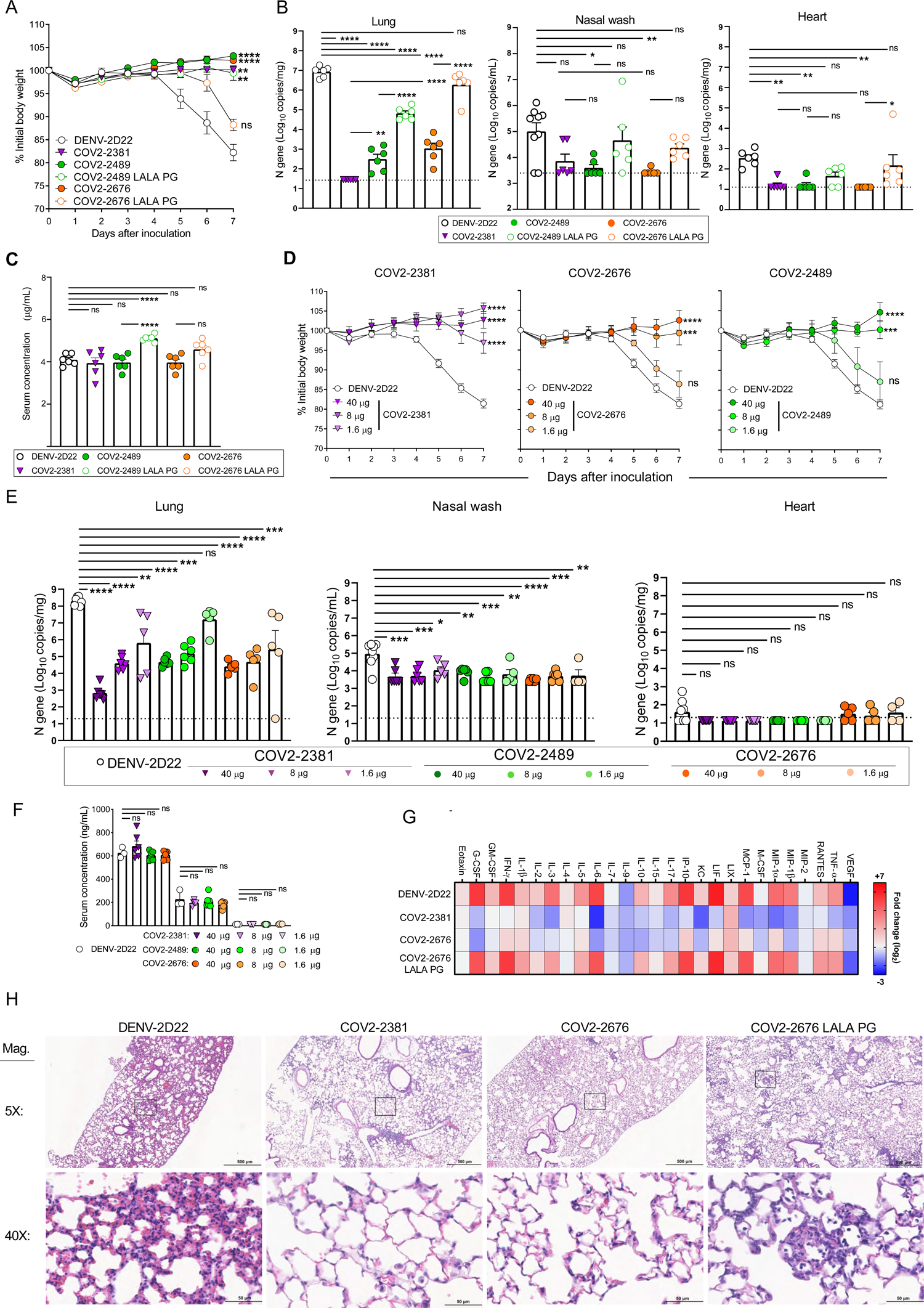
Prophylaxis with COV2-2676 or COV2-2489 confers protection against SARS-CoV-2 in mice. Eight to nine week-old male and female K18 hACE2 transgenic mice were administered by intraperitoneal (i.p.) injection 200, 40, 8 or 1.6 μg of COV2-2676, COV2-2489, COV2-2676 LALA-PG, COV2-2489 LALA-PG, COV2-2381 (a positive control) or DENV-2D22, an isotype control, mAb a day before virus inoculation (D-1). One day later, mice were inoculated intranasally (i.n.) with 10^3^ PFU of SARS-CoV-2. **A.** Body weight change of mice over time. Data shows the mean ± S.E.M. compared to the isotype control mAb for two independent experiments (n = 6 to 11 for each experimental group; one-way ANOVA with Dunnett’s post hoc test of area under the curve from 4 to 7 dpi: ns, not significant, ^∗∗^p < 0.01,^∗∗∗∗^p < 0.000 1). **B.** Tissues were harvested at 7 dpi from a subset of mice in **A**. Viral burden in the lung, nasal wash, and heart was assessed by qRT-PCR of the N gene. Data shows the mean ± S.E.M. compared between all groups for two independent experiments (n = 6 for each experimental group; one-way ANOVA with Tukey’s post hoc test: ns, not significant, ^∗^p < 0.05, ^∗∗^p < 0.01, ^∗∗∗^p < 0.001, ^∗∗∗∗^p < 0.0001). Dashed line represents limit of detection of assay. **C.** Serum concentration (ng/ml) of human mAbs at the time of SARS-CoV-2 infection of the mice in **B**. Data shows the mean ± S.E.M. compared between all groups for two independent experiments (n = 6 for each experimental group; one-way ANOVA with Tukey’s post hoc test: ns, not significant, ^∗∗∗∗^p < 0.0001). **D.** Body weight change of mice over time. Data shows the mean ± S.E.M. compared to isotype control mAb for two independent experiments (n = 4 to 8 for each experimental group; one-way ANOVA with Dunnett’s post hoc test of area under the curve from 4 to 7 dpi: ns, not significant, ^∗∗∗^p < 0.001, ^∗∗∗∗^p < 0.0001). **E.** Tissues were harvested at 7 dpi from mice in **D**. Viral burden in the lung, nasal wash, and heart was assessed by qRT-PCR of the N gene. Data shows the mean ± S.E.M. compared between all groups for two independent experiments (n = 4 to 8 for each experimental group; one-way ANOVA with Tukey’s post hoc test: ns, not significant, ^∗^p < 0.05, ^∗∗^p < 0.01, ^∗∗∗^p < 0.001, ^∗∗∗∗^p < 0.0001). **F.** Serum concentration (ng/mL) of human mAbs at the time of SARS-CoV-2 infection of the mice in **D**. Data shows the mean ± S.E.M. compared to isotype control mAb at various doses of mAb for two independent experiments (n = 6; one-way ANOVA with Dunnett’s post hoc test: ns, not significant). **G.** Heat map of cytokine and chemokine levels in lung tissue homogenates harvested in **B** as measured by a multiplex platform. Log_2_ fold change compared to lungs from mock-infected animals was plotted in the corresponding heat map (associated statistics are reported in **Fig S6A**). **H.** Hematoxylin and eosin staining of lung sections harvested at 7 dpi from mice in **B**. Images are low (top; scale bars, 500 μm) and high power (bottom; scale bars, 50 μm). Representative images are shown from two independent experiments (n = 3).

We next assessed the therapeutic efficacy of monotherapy with COV2-2676 or COV-2489 in a post-exposure setting. Mice treated with 200 μg of COV2-2676 or COV2-2489 one day after SARS-CoV-2 inoculation maintained weight, in a manner similar to COV2-2381 treatment and in contrast to the isotype control mAb, DENV-2D22 (**Fig 5A**). Therapy with COV2-2676 or COV2-2489 also decreased viral burden at 7 dpi in the upper and lower respiratory tract and heart, compared to DENV-2D22, in a manner similar to COV2-2381 (**Fig 5B**). Animals treated with COV2-2676 also had lower levels of multiple cytokines and chemokines in lung homogenates, similar to COV2-2381 and in contrast to DENV-2D22 (**Fig 5C** and **S5B**). Pathological analysis also showed less immune cell infiltration and airspace consolidation after COV2-2676 or COV2-2381 therapy compared to DENV-2D22 treatment (**Fig 5D**).

**Figure 5.**
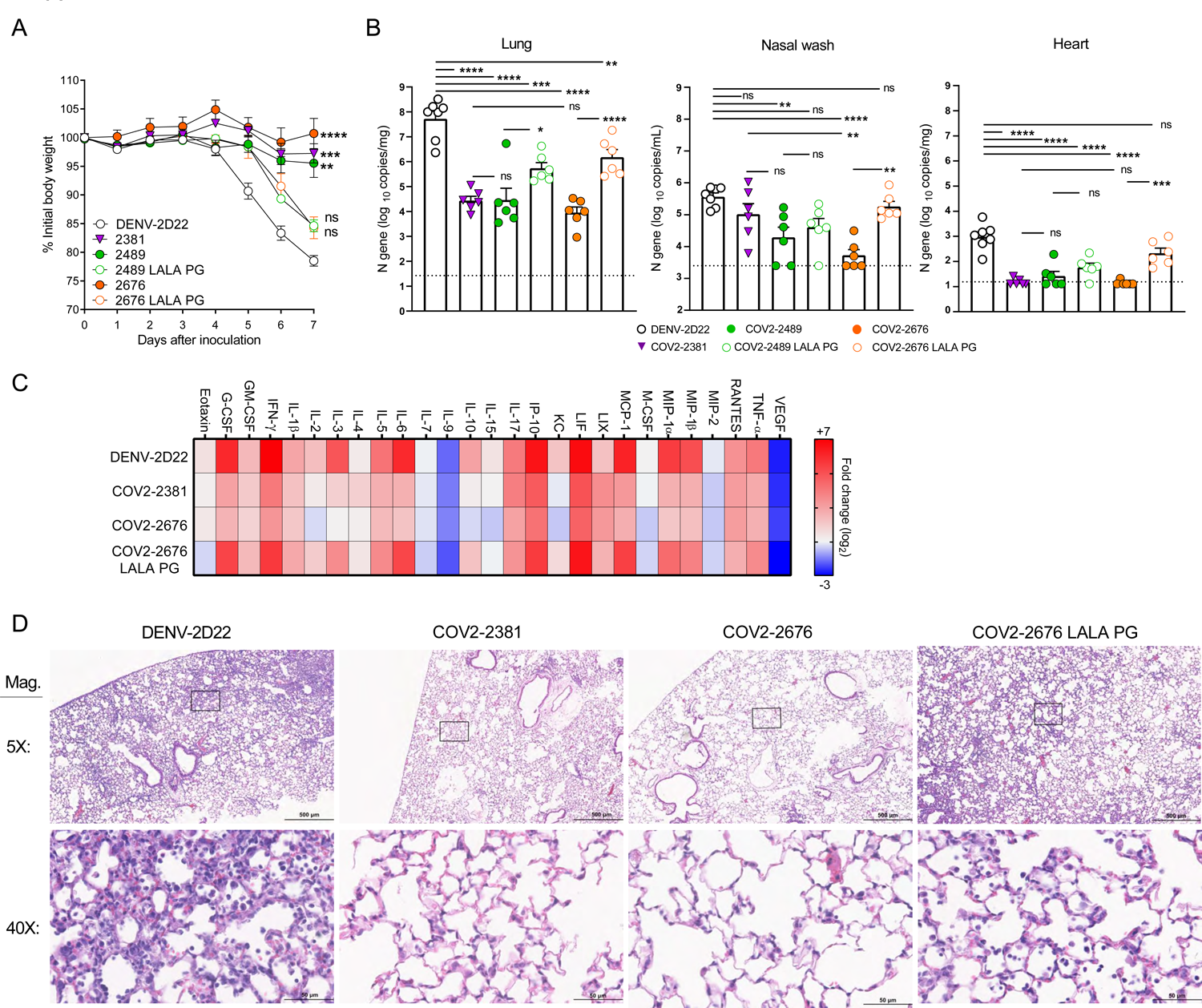
Therapeutic activity of COV2-2676 or COV-2489 after SARS-CoV-2 challenge. Eight to nine week-old male and female K18 hACE2 transgenic mice were inoculated with 10^3^ PFU of SARS-COV2. One day later (D+1), mice were given an i.p. administration of 200 μg of COV2-2676, COV2-2489, COV2-2676 LALA PG, COV2-2489 LALA PG, COV2-2381 (a positive control) or DENV-2D22, an isotype control, mAb. **A.** Body weight change of mice over time. Data shows the mean ± S.E.M. compared to isotype control mAb for two independent experiments (n = 6 to 11 for each experimental group; one-way ANOVA with Dunnett’s post hoc test of area under the curve from 4 to 7 dpi: ns, not significant, ∗∗p < 0.01, ∗∗∗p <0.001, ∗∗∗∗p < 0.0001). **B.** Tissues were harvested at 7 dpi from a subset of mice in **A**. Viral burden in the lung, nasal wash, and heart was assessed by qRT-PCR of the N gene. Data shows the mean ± S.E.M. compared between all groups for two independent experiments (n = 6 to 7 for each experimental group; one-way ANOVA with Tukey’s post hoc test: ns, not significant, ∗p < 0.05, ∗∗p < 0.01, ∗∗∗p < 0.001, ∗∗∗∗p < 0.0001). **C.** Heat map of cytokine and chemokine levels in lung homogenates harvested in **B** as measured by a multiplex platform. Log_2_ fold change compared to lungs from mock-infected animals was plotted in the corresponding heat map (associated statistics are reported in **Fig S6B**). **D.** Hematoxylin and eosin staining of lung sections harvested at 7 dpi from mice in **B**. Images are low (top; scale bars, 500 μm) and high power (bottom; scale bars, 50 μm). Representative images are shown from two independent experiments (n = 3).

### Fc effector functions contribute to optimal protection by COV2-2676 or COV-2489

Because the NTD mAbs bound avidly to the surface of SARS-CoV-2-infected cells (**Fig 3A** and **S4A**), we speculated that part of their protective activity might be mediated by effector functions through Fc engagement of C1q or FcγRs that could promote clearance. To test this hypothesis, we engineered LALA-PG mutations into the Fc region of these human IgG1 mAbs to abrogate interaction of the Fc region with FcγRs and complement proteins (Lund et al., 1991; Wines et al., 2000). We confirmed that the LALA-PG mutations in COV2-2676 and COV2-2489 did not affect S protein binding (**Fig S4B**) or neutralizing activity (**Fig S4C**).

We then evaluated if the LALA-PG Fc variants lost protective activity *in vivo*. Despite similar concentrations in serum at the time of SARS-CoV-2 infection (**Fig 4C**), increased weight loss and viral burden, as well as greater lung inflammation and pathology were observed in animals pre-treated with COV2-2676 LALA-PG Fc or COV2-2489 LALA-PG Fc variant IgGs compared to the intact parental mAbs (**Fig 4A, B, E, G, and H**). This pattern also was seen with therapy initiated one day after SARS-CoV-2 infection using COV2-2676 LALA-PG Fc or COV2-2489 LALA-PG Fc variants (**Fig 5A-D**). Overall, these findings suggest that Fc effector function activities contribute to the protection conferred by each of the NTD mAbs.

The combination of an RBD-specific neutralizing mAb and an NTD mAb in a cocktail would confer equivalent or better levels of protection, since binding to distinct antigenic sites might mitigate the risk of selection of antibody escape variants. Thus, mAb cocktails that include components binding to different epitopes on S protein offer higher resistance to escape mutations (Baum et al., 2020b; Greaney et al., 2021) and protect animals from SARS-CoV-2 challenge (Baum et al., 2020a; Zost et al., 2020a). Initially, to test this approach, we used VSV expressing escape variants of the SARS-CoV-2 S protein that were resistant to neutralization by RBD-specific antibodies (COV-2479 or COV2-2130) (Greaney et al., 2021) and the NTD-specific antibodies described here (COV2-2676 and COV2-2489). As expected, NTD-reactive mAbs neutralized the RBD mAb escape mutant viruses, and RBD-reactive mAbs neutralized the NTD mAb escape mutant viruses (**Fig 6A**).

**Figure 6.**
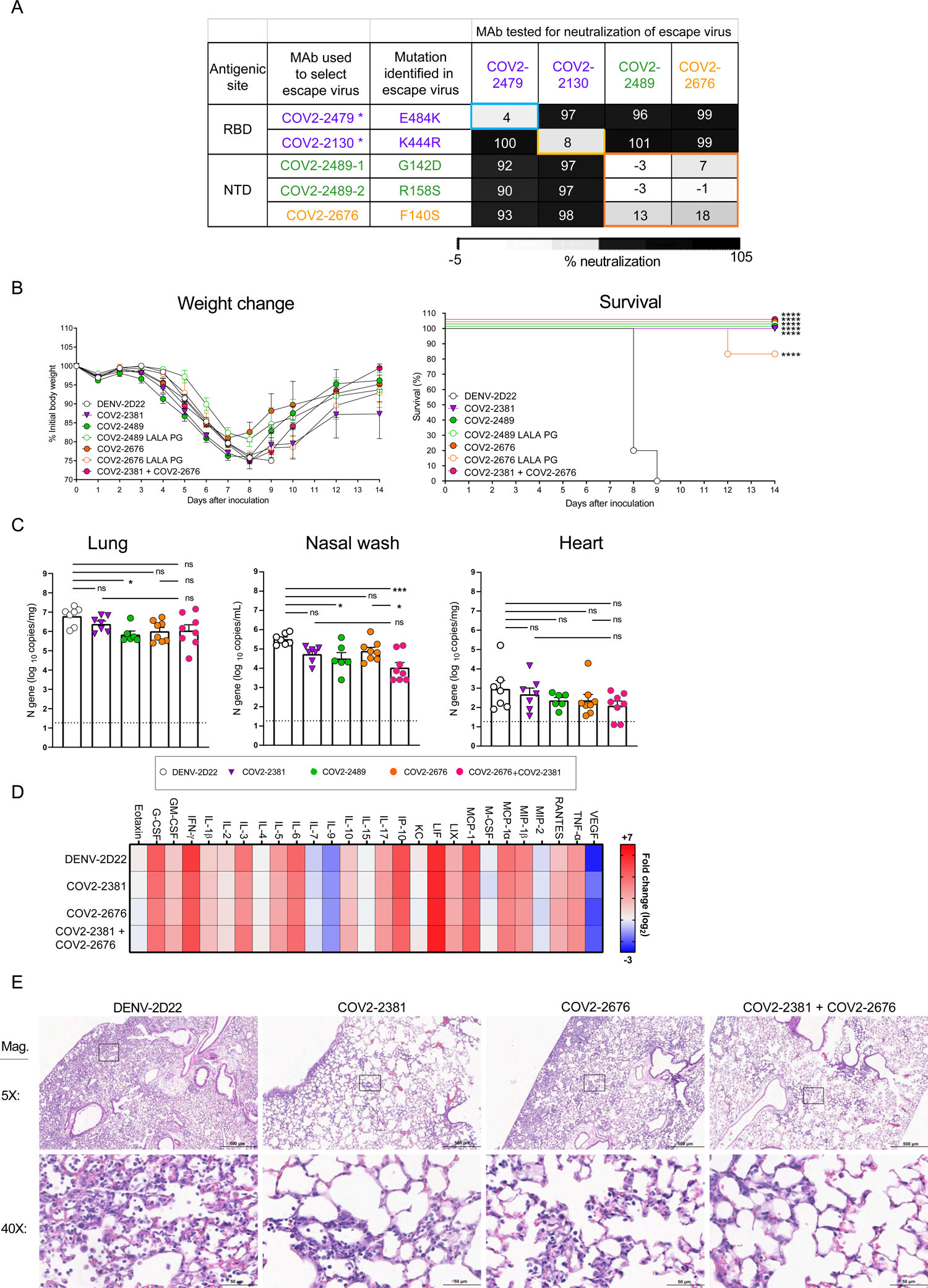
Rationale for design of an antibody cocktail targeting both RBD and NTD. **A.** Neutralization matrix of RBD mAbs (COV2-2479 and COV2-2130 shown in purple) and NTD mAb (COV2-2676 in orange and COV2-2489 in green) escape viruses. Black = full neutralization, grey = partial neutralization, white = no neutralization & * indicates escape viruses that were isolated in previously published study. Eight to nine week-old male or female K18 hACE2 transgenic mice were inoculated intranasally with 10^3^ PFU of SARS-CoV2. Two days later (D+2), mice were given an i.p. administration of 200 μg of COV2-2676, COV2-2489, COV2-2676 LALA-PG, COV2-2489 LALA-PG, COV2-2381 (a positive control) or DENV-2D22, an isotype control, mAb for monotherapy or 100 µg each of COV2-2381 and COV2-2676 for combination therapy. **B.** (Left panel) Body weight change of mice over time. Data shows the mean ± S.E.M. compared to isotype control mAb for two independent experiments (n = 6 to 14 for each experimental group: one-way ANOVA with Dunnett’s post hoc test of area under the curve from 4 to 7 dpi. Significant differences were not detected. (Right panel) Percent survival of mice over time. Survival was compared to isotype control for two independent experiments (n = 6 to 14 for each experimental group; log-rank Mantel-Cox test; ^∗∗∗∗^p < 0.0001). **C.** Tissues were harvested at 7 dpi mice from a subset of mice in **B**. Viral burden in the lung, nasal wash, and heart was assessed by qRT-PCR of the N gene. Data shows the mean ± S.E.M. compared between all groups for two independent experiments: n = 6 to 8 for each experimental group; one-way ANOVA with Tukey’s post hoc test: ns, not significant, ^∗^p < 0.05, ^∗∗∗^p <0.001). **D.** Heat map of cytokine and chemokine levels in lung homogenates harvested in **C** as measured by a multiplex platform. Log_2_ fold change compared to lungs from mock-infected animals was plotted in the corresponding heat map (associated statistics are reported in **Fig S6C**.) **E.** Hematoxylin and eosin staining of lung sections harvested at 7 dpi from mice in **B**. Images are low (top; scale bars, 500 μm) and high power (bottom; scale bars, 50 μm). Representative images are shown from two independent experiments (n = 3).

We next treated mice two days after inoculation with SARS-CoV-2 with individual mAbs or a combination of COV2-2381 (RBD-specific) and COV2-2676 (NTD-specific) mAbs. Mice treated with 200 μg of COV2-2676, COV2-2489 or COV2-2831 two days after SARS-CoV-2 inoculation showed weight loss, viral burden, and inflammatory mediator profiles that were similar to DENV-2D22-treated mice (**Fig 6B-D**), consistent with a limited therapeutic window for effective treatment in this stringent transgenic hACE2 mouse model. While mice were not protected from the initial phase of weight loss, viral infection in the lower respiratory tract or heart, or lung inflammation (**Fig 6B-D, S5C),** treatment with mAb monotherapy or the cocktail resulted in weight gain beginning approximately one week after infection and survival through two weeks after infection, and this recovery was associated with decreased nasal wash titers and reduced lung pathology (**Fig 6B-D**). In comparison, all mice treated with the isotype control mAb DENV-2D22 had severe lung pathology and succumbed to infection between eight and nine days after inoculation. These results suggested that diversification of neutralizing antibody responses after natural infection or vaccination (*e.g*., targeting of vulnerable RBD and NTD sites of the spike) could be beneficial for protective immunity, with a potential implication for therapeutic cocktail design.

## DISCUSSION

Several research groups have identified RBD-specific mAbs using B cells from SARS-CoV-2 convalescent individuals, with a number of these mAbs now in Phase III clinical trials (Baum et al., 2020a; Chen et al., 2020a; Jones et al., 2020) and two mAb products having obtained Emergency Use Authorization from the U.S. Food and Drug Administration. Due to the large number of natural variant viruses with polymorphisms in the RBD that are emerging, identification of neutralizing and protective human mAbs that bind to other antigenic sites on the S protein is warranted. In a large (n =389) panel of mAbs we isolated with reactivity to the SARS-CoV-2 S protein, the majority of mAbs bound to S protein but not to the RBD (Zost et al., 2020b). We found that a small subset of non-RBD antibodies that recognized the NTD could neutralize SARS-CoV-2 infection and confer protection in a stringent hACE2 transgenic mouse model of disease.

Our studies highlight the promising functional activities of NTD-specific mAbs. Although these mAbs did not bind as avidly to recombinant S6_Pecto_ protein as did the most potent RBD-reactive mAbs we isolated, they neutralized SARS-CoV-2 with potencies comparable to many RBD-targeting antibodies (Zost et al., 2020a; Zost et al., 2020b). Very few NTD-targeting neutralizing antibodies have been reported to date. Sequence comparison of recently published NTD-reactive SARS-CoV-2 mAbs 4A8 (Chi et al., 2020) and 4-8 (Liu et al., 2020a) revealed that our mAbs are genetically distinct and not members of a public clonotype matching those mAbs (**Table S1**).

Most neutralizing SARS-CoV-2 mAbs block RBD interactions with ACE2 (Brouwer et al., 2020; Cao et al., 2020; Ju et al., 2020; Liu et al., 2020a; Rogers et al., 2020; Seydoux et al., 2020; Wec et al., 2020; Wu et al., 2020; Zost et al., 2020a; Zost et al., 2020b). However, the mechanism of neutralization of NTD-reactive antibodies appears complex. Our data suggest that the anti-NTD antibodies COV2-2676 and COV2-2489 inhibit at a post-attachment phase of infection and may block subsequent virus entry or fusion steps. In cells in which anti-RBD and anti-NTD antibodies do not appear to block virus attachment (*i.e*., conventional Vero cells), both groups inhibited at a post-attachment step. Virus attachment to cells can differ in diverse cells, likely because of the capacity of SARS-CoV-2 to use multiple attachment factors including heparan sulfates, hACE2, TAM receptors, and possibly other molecules (Cantuti-Castelvetri et al., 2020; Chen et al., 2020b; Chiodo et al., 2020; Clausen et al., 2020; Gao and Zhang, 2020). The mechanism of attachment in Vero (monkey kidney) cells without human ACE2 transfection is uncertain; attachment might be mediated by monkey ACE2 although this has not been formally demonstrated. It is possible that the NTD antibodies block entry by indirectly interfering with ACE2 binding through steric interference of the Fc region. In analogous experiments with MERS-CoV, structural studies presented showed that the MERS-CoV mAb 7D10 could bind to the NTD of S protein of MERS-CoV and inhibit RBD-DPP4 interaction through its light chain, which was close to the RBD (Zhou et al., 2019). We tested that model here but found that neither COV2-2676 nor COV2-2489 blocked ACE2 interaction with RBD in soluble protein or in cell-surface-display assays. Antibody fragment studies using Fab, F(ab′)_2_ and IgG forms of COV2-2676 and COV2-2489 showed that the Fab forms lost neutralizing activity, possibly suggesting some blocking occurs indirectly by steric effects of the projecting Fc region into the direction of the RBD as in the case for the MERS-CoV mAB 7D10 (Zhou et al., 2019). However, we did not obtain direct evidence for this mechanism. In cells expressing high hACE2 levels, anti-RBD antibodies partially inhibited attachment of virus, but anti-NTD antibodies did not.

To define the fine specificity of this protective epitope, we performed saturation alanine-scanning mutagenesis studies and identified critical residues for binding of COV2-2676 or COV2-2489 mAbs in the NTD. We showed that SARS-CoV-2 S protein variants with mutations at F140S and G142D, or R158S in the NTD conferred resistance to COV2-2676 or COV2-2489, respectively, using a chimeric replication-competent VSV expressing the SARS-CoV-2 S protein. Consistent with these data, Weisblum *et al*. identified an NTD epitope (residues 148 to 151), and mutations in this epitope enabled escape from neutralizing antibody activity in a donor plasma specimen (Weisblum et al., 2020).

We first tested the prophylactic efficacy of COV2-2676 or COV2-2489 in a well-defined model of SARS-CoV-2 infection in hACE2-expressing transgenic mice (Winkler *et al*., 2020). Mice pre-treated with COV2-2676 or COV2-2489 mAbs exhibited substantially lower viral titers than mice treated with an isotype-control mAb and offered complete protection against death in this model. We also assessed the therapeutic potential of these mAbs in the same mouse model. Each of the mAbs mediated a therapeutic effect when administered after infection. Although several protection studies of anti-SARS-CoV-2 mAbs have been published, all of these experiments used mAbs directed against the RBD (Baum et al., 2020a; Chen et al., 2020a; Hassan et al., 2020; Wang et al., 2020; Zost et al., 2020a). There is only limited precedence for *in vivo* protection against SARS-CoV-2 infection or disease mediated by mAbs that react to S protein regions outside the RBD (McCallum et al., 2021; Voss et al., 2020; Zhang et al., 2020).

IgG interactions with Fcγ receptors and complement contribute to antibody-dependent viral clearance during many infections, including HIV-1, Ebola, influenza, and chikungunya viruses (DiLillo et al., 2014; Fox et al., 2019; Halper-Stromberg et al., 2014; Lu et al., 2016). MAbs against SARS-CoV-1 and SARS-CoV-2 also can confer protective effects in part through through Fc-mediated effector functions (Atyeo et al., 2020; Pinto et al., 2020; Schafer et al., 2021). To test whether this was the case with our mAbs, we generated COV2-2676 and 2489 LALA-PG variant Fc versions. Indeed, Fc-mediated activity for NTD-specific mAbs did contribute to protection in the models we tested. Animals in the LALA-PG mAb-treated mice groups had increased viral burden and lost weight, but still survived viral challenge, suggesting involvement of Fc-effector function dependent and independent mechanisms in the protection *in vivo* conferred by anti-NTD mAbs.

SARS-CoV-2 continues to evolve, and its capacity to escape the activity of neutralizing antibodies in current clinical development is not fully understood. The emergence of mutations at E484K in South African strains is concerning, because this change can impact the neutralizing activity of many RBD-specific mAbs and immune serum generated from convalescent subjects (Avanzato et al., 2020; Greaney et al., 2021; Liu et al., 2020b; Oude Munnink et al., 2021; Piccoli et al., 2020; Rambaut et al., 2020; Starr et al., 2020). Moreover, the administration of convalescent plasma with suboptimal levels of neutralizing antibodies might increase resistance in circulating SARS-CoV-2 populations (Bloch, 2020). Furthermore, candidate vaccines that include only RBD antigens lack the ability to induce NTD-reactive neutralizing antibodies (Laczko et al., 2020; Mulligan et al., 2020; Smith et al., 2020; Yu et al., 2020). Antibody therapy or vaccine approaches that target additional antigenic sites may limit escape and prevent compromising of vaccine- or natural infection-induced immunity. A combination of RBD- and NTD-reactive neutralizing mAbs may offer an attractive alternative approach to treatments that target only RBD. The data presented here describing neutralization escape viruses suggests the potential utility of using mAb cocktails to avoid selection of neutralization resistant viruses. Moreover, we establish that a combination of RBD- and NTD-reactive neutralizing mAbs can be used as an effective therapeutic cocktail *in vivo*. Overall, the results presented here provide compelling evidence that some NTD-targeting mAbs can inhibit SARS-CoV-2 infection efficiently *in vitro* and *in vivo*, using both neutralizing and Fc-mediated effector function activities.

## ACKNOWLEDGEMENTS

We thank Ryan Irving at Vanderbilt University Medical Center for laboratory management support and Merissa Mayo and Norma Suazo Galeano for human subjects support. EM data collections were conducted at the Center for Structural Biology Cryo-EM Facility at Vanderbilt University. This work was supported by the NIAID/NIH grants T32 AI095202 (S.J.Z.), T32 AI007163 (E. S. W.), and R01 AI157155 (M.S.D. and J.E.C.), HHSN contracts 75N93019C00074 (J.E.C.) and 75N93019C00073 (B.J.D.), DARPA HR0011-18-2-0001 (J.E.C.); the Dolly Parton COVID-19 Research Fund at Vanderbilt (J.E.C.); and Fast Grants, Mercatus Center, George Mason University (J.E.C.). J.E.C. is a recipient of the 2019 Future Insight Prize from Merck KGaA. The content is solely the responsibility of the authors and does not represent the official views of the U.S. government or other sponsors.

## AUTHOR CONTRIBUTIONS

Conceived of the project: N.S., P.G., S.J.Z., R.H.C., L.B.T., M.S.D., J.E.C.; Obtained funding: M.S.D. and J.E.C. Performed laboratory experiments: N.S., S.S., P.G., E.S.W., E.B., S.J.Z., L.V., R.S.N., R.E.S., S.P.K., M.E.F., L.B.T. Performed computational work: E.C.C. Supervised research: R.H.C., L.B.T., E.D., B.D., M.S.D., J.E.C. Wrote the first draft of the paper: N.S., P.G., S.S., L.B.T., M.S.D., J.E.C.; All authors reviewed and approved the final manuscript.

## STAR METHODS

### RESOURCE AVAILABILITY

#### LEAD CONTACT

Further information and requests for resources and reagents should be directed to and will be fulfilled by the Lead Contact, James E. Crowe, Jr.

#### MATERIALS AVAILABILITY

Materials described in this paper are available for distribution for nonprofit use using templated documents from Association of University Technology Managers “Toolkit MTAs”, available at: https://autm.net/surveys-and-tools/agreements/material-transfer-agreements/mta-toolkit.

#### DATA AND CODE AVAILABILITY

All data needed to evaluate the conclusions in the paper are present in the paper or the Supplemental Information. The antibodies in this study are available by Material Transfer Agreement with Vanderbilt University Medical Center.

### EXPERIMENTAL MODEL AND SUBJECT DETAILS

#### Antibodies

The human antibodies studied in this paper were isolated from blood samples from two individuals in North America with previous laboratory-confirmed symptomatic SARS-CoV-2 infection that was acquired in China. The original clinical studies to obtain specimens after written informed consent were previously described (Zost et al., 2020b) and approved by the Institutional Review Board of Vanderbilt University Medical Center, the Institutional Review Board of the University of Washington and the Research Ethics Board of the University of Toronto. The individuals (a 56-year-old male and a 56-year-old female) are a married couple and residents of Wuhan, China who travelled to Toronto, Canada, where PBMCs were obtained by leukopheresis 50 days after symptom onset. The antibodies were isolated using diverse tools for isolation and cloning of single antigen-specific B cells and the antibody variable genes that encode mAbs (Zost et al., 2020b).

#### Cell lines

Vero E6 (ATCC, CRL-1586), Vero (ATCC, CCL-81), HEK293 (ATCC, CRL-1573) and HEK293T (ATCC, CRL-3216) cells were maintained at 37°C in 5% CO2 in Dulbecco’s minimal essential medium (DMEM) containing 10% (v/v) heat-inactivated fetal bovine serum (FBS), 10 mM HEPES pH 7.3, 1 mM sodium pyruvate, 1× non-essential amino acids and 100 U/mL of penicillin– streptomycin. Vero-furin cells were obtained from T. Pierson (NIAID, NIH) and have been described previously (Mukherjee et al., 2016). Vero-hACE2-TMPRSS2 cells were a gift of A. Creanga and B. Graham (Vaccine Research Center, NIH). FreeStyle 293F cells (Thermo Fisher Scientific, R79007) were maintained at 37°C in 8% CO2. Expi293F cells (Thermo Fisher Scientific, A1452) were maintained at 37°C in 8% CO2 in Expi293F Expression Medium (Thermo Fisher Scientific, A1435102). ExpiCHO cells (Thermo Fisher Scientific, A29127) were maintained at 37°C in 8% CO2 in ExpiCHO Expression Medium (Thermo Fisher Scientific, A2910002). Authentication analysis was not performed for the cell lines used. Mycoplasma testing of Expi293F and ExpiCHO cultures was performed on a monthly basis using a PCR-based mycoplasma detection kit (ATCC, 30-1012K).

#### Viruses

SARS-CoV-2 strain 2019 n-CoV/USA_WA1/2020 was obtained from the Centers for Disease Control and Prevention (a gift from N. Thornburg). Virus was passaged in Vero CCL81 cells and titrated by plaque assay on Vero E6 cell culture monolayers as previously described (Case et al., 2020a). The generation of a replication-competent VSV expressing SARS-CoV-2 S protein with a 21 amino-acid C-terminal deletion that replaces the VSV G protein (VSV-SARS-CoV-2) was described previously (Case et al., 2020b). The S protein-expressing VSV virus was propagated in MA104 cell culture monolayers (African green monkey, ATCC CRL-2378.1) as described previously (Case *et al*., 2020), and viral stocks were titrated on Vero E6 cell monolayer cultures. VSV plaques were visualized using neutral red staining. All work with infectious SARS-CoV-2 was performed in Institutional Biosafety Committee approved BSL3 and A-BSL3 facilities at Washington University School of Medicine using appropriate positive pressure air respirators and protective equipment.

#### Mouse models

Animal studies were carried out in accordance with the recommendations in the Guide for the Care and Use of Laboratory Animals of the National Institutes of Health. The protocols were approved by the Institutional Animal Care and Use Committee at the Washington University School of Medicine (assurance number A3381–01). Virus inoculations were performed under anesthesia that was induced and maintained with ketamine hydrochloride and xylazine, and all efforts were made to minimize animal suffering. Heterozygous K18-hACE c57BL/6J mice (strain: 2B6.Cg-Tg(K18-ACE2)2Prlmn/J) were obtained from Jackson Laboratory (034860). Eight to nine week-old mice of both sexes were inoculated with 10^3^ PFU of SARS-CoV-2 by an intranasal route.

## METHOD DETAILS

### Recombinant antigens and proteins

A gene encoding the ectodomain of a pre-fusion conformation-stabilized SARS-CoV-2 S protein ectodomain (S6_Pecto_) (Hsieh et al., 2020) was synthesized and cloned into a DNA plasmid expression vector for mammalian cells. A similarly designed S protein antigen with two prolines and removal of the furin cleavage site for stabilization of the prefusion form of S (S2_Pecto_) was reported previously (Wrapp et al., 2020). In brief, this gene includes the ectodomain of SARS-CoV-2 (to residue 1,208), a T4 fibritin trimerization domain, an AviTag site-specific biotinylation sequence and a C-terminal 8× His tag. To stabilize the construct in the pre-fusion conformation, we included substitutions F817P, A892P, A899P, A942P, K986P and V987P and mutated the furin cleavage site at residues 682–685 from RRAR to ASVG. The recombinant S6_Pecto_ protein was isolated by metal affinity chromatography on HisTrap Excel columns (Cytiva), and protein preparations were purified further by size-exclusion chromatography on a Superose 6 Increase 10/300 column (Cytiva). The presence of trimeric, pre-fusion conformation S protein was verified by negative-stain electron microscopy (Zost et al., 2020b). For electron microscopy with S protein and Fabs, we expressed a variant of S6_Pecto_ lacking an AviTag but containing a C-terminal Twin-Strep-tag, similar to that described previously (Zost et al., 2020b). Expressed protein was isolated by metal affinity chromatography on HisTrap Excel columns (Cytiva), followed by further purification on a StrepTrap HP column (Cytiva) and size-exclusion chromatography on TSKgel G4000SWXL (TOSOH). To express the RBD subdomain of the SARS-CoV-2 S protein, a synthetic DNA (Twist Bioscience) encoding residues 319–541 was cloned into a mammalian expression vector downstream of an IL-2 signal peptide and upstream of a thrombin cleavage site, an AviTag and a 6× His tag. Recombinant SARS-CoV-2 S NTD protein was kindly provided by P. McTamney, K. Ren and A. Barnes (AstraZeneca). Purified proteins were analyzed by SDS–PAGE to ensure purity and appropriate molecular weights.

### MAb production and purification

Sequences of mAbs that had been synthesized (Twist Bioscience) and cloned into an IgG1 monocistronic expression vector (designated as pTwist-mCis_G1) or Fab expression vector (designated as pTwist-mCis_FAB) were used for production in mammalian cell culture. This vector contains an enhanced 2A sequence and GSG linker that allows the simultaneous expression of mAb heavy and light chain genes from a single construct upon transfection (Chng et al., 2015). For antibody production, we performed transfection of ExpiCHO cell cultures using the Gibco ExpiCHO Expression System as described by the vendor. IgG molecules were purified from culture supernatants using HiTrap MabSelect SuRe (Cytiva) on a 24-column parallel protein chromatography system (Protein BioSolutions). Fab proteins were purified using CaptureSelect column (Thermo Fisher Scientific). Purified antibodies were buffer-exchanged into PBS, concentrated using Amicon Ultra-4 50-kDa (IgG) or 30 kDa (Fab) centrifugal filter units (Millipore Sigma) and stored at 4°C until use. F(ab′)_2_ fragments were generated after cleavage of IgG with IdeS protease (Promega) and then purified using TALON metal affinity resin (Takara) to remove the enzyme and protein A agarose (Pierce) to remove the Fc fragment. Purified mAbs were tested routinely for endotoxin levels (found to be less than 30 EU per mg IgG). Endotoxin testing was performed using the PTS201F cartridge (Charles River), with a sensitivity range from 10 to 0.1 EU per mL, and an Endosafe Nexgen-MCS instrument (Charles River).

### ELISA binding assays

Wells of 96-well microtiter plates were coated with purified recombinant SARS-CoV-2 S6_Pecto_, SARS-CoV-2 S NTD, or SARS-CoV-2 RBS protein at 4 °C overnight. Plates were blocked with 2% non-fat dry milk and 2% normal goat serum in Dulbecco’s phosphate-buffered saline (DPBS) containing 0.05% Tween-20 (DPBS-T) for 1 h. The bound antibodies were detected using goat anti-human IgG conjugated with horseradish peroxidase (HRP) (Southern Biotech, cat. 2040-05, lot B3919-XD29, 1:5,000 dilution) and a 3,3′,5,5′-tetramethylbenzidine (TMB) substrate (Thermo Fisher Scientific). Color development was monitored, 1M HCl was added to stop the reaction, and the absorbance was measured at 450 nm using a spectrophotometer (Biotek). For dose–response assays, serial dilutions of purified mAbs were applied to the wells in triplicate, and antibody binding was detected as detailed above. EC50 values for binding were determined using Prism v.8.0 software (GraphPad) after log transformation of the mAb concentration using sigmoidal dose–response nonlinear regression analysis.

### Focus reduction neutralization test (FRNT)

Serial dilutions of mAbs were incubated with 10^2^ FFU of SARS-CoV-2 for 1 h at 37 °C. The antibody-virus complexes were added to Vero E6 cell-culture monolayers in 96-well plates for 1 h at 37 °C. Cells then were overlaid with 1% (w/v) methylcellulose in minimum essential medium (MEM) supplemented to contain 2% heat-inactivated FBS. Plates were fixed 30 h later by removing overlays and fixed with 4% paraformaldehyde (PFA) in PBS for 20 min at room temperature. The plates were incubated sequentially with 1 μg/mL of rCR3022 anti-S antibody or an murine anti-SARS-COV-2 mAb, SARS2-16 (hybridoma supernatant diluted 1:6,000 to a final concentration of ∼20 ng/mL) and then HRP-conjugated goat anti-human IgG (Sigma-Aldrich, A6029) in PBS supplemented with 0.1% (w/v) saponin (Sigma) and 0.1% BSA. SARS-CoV-2-infected cell foci were visualized using TrueBlue peroxidase substrate (KPL) and quantitated on an ImmunoSpot 5.0.37 Macro Analyzer (Cellular Technologies). IC_50_ values were determined by nonlinear regression analysis (with a variable slope) using Prism software.

### Real-time cell analysis (RTCA) neutralization assay

To determine neutralizing activity of IgG, Fab, or F(ab′)_2_ proteins, we used real-time cell analysis (RTCA) assay on an xCELLigence RTCA MP Analyzer (ACEA Biosciences Inc.) that measures virus-induced cytopathic effect (CPE) (Gilchuk et al., 2020a; Zost et al., 2020b). Briefly, 50 μL of cell culture medium (DMEM supplemented with 2% FBS) was added to each well of a 96-well E-plate using a ViaFlo384 liquid handler (Integra Biosciences) to obtain background reading. A suspension of 18,000 Vero-E6 cells in 50 μL of cell culture medium was seeded in each well, and the plate was placed on the analyzer. Measurements were taken automatically every 15 min, and the sensograms were visualized using RTCA software version 2.1.0 (ACEA Biosciences Inc). VSV-SARS-CoV-2 (0.01 MOI, ∼120 PFU per well) was mixed 1:1 with a dilution of mAb in a total volume of 100 μL using DMEM supplemented with 2% FBS as a diluent and incubated for 1 h at 37°C in 5% CO2. At 16 h after seeding the cells, the virus-mAb mixtures were added in replicates to the cells in 96-well E-plates. Triplicate wells containing virus only (maximal CPE in the absence of mAb) and wells containing only Vero cells in medium (no-CPE wells) were included as controls. Plates were measured continuously (every 15 min) for 48 h to assess virus neutralization. Normalized cellular index (CI) values at the endpoint (48 h after incubation with the virus) were determined using the RTCA software version 2.1.0 (ACEA Biosciences Inc.). Results are expressed as percent neutralization in a presence of respective mAb relative to control wells with no CPE minus CI values from control wells with maximum CPE. RTCA IC_50_ values were determined by nonlinear regression analysis using Prism software.

### Pre- and post-attachment neutralization assays

For pre-attachment assays, serial dilutions of mAbs were prepared at 4°C in Dulbecco’s modified Eagle medium (DMEM) with 2% FBS and preincubated with 10^2^ FFU of SARS-CoV-2 for 1 h at 4°C. MAb-virus complexes were added to a monolayer of Vero cells for 1 h at 4°C. Virus was allowed to internalize during a 37°C incubation for 30 min. Cells were overlaid with 1% (wt/vol) methylcellulose in MEM. For post-attachment assays, 2 x 10^2^ FFU of SARS-CoV-2 was adsorbed onto a monolayer of Vero cells for 1 h at 4°C. After removal of unbound virus, cells were washed twice with cold DMEM, followed by the addition of serial dilutions of MAbs in cold DMEM. Virus-adsorbed cells were incubated with mAd dilutions for 1 h at 4°C. Virus then was allowed to internalize for 30 min at 37°C, and subsequently cells were overlaid with methylcellulose as described above. Thirty hours later, the plates were fixed with 4% PFA and analyzed for antigen-specific foci as described above for FRNTs.

### Attachment inhibition assay

SARS-COV-2 was incubated with mAbs at the specified concentration for 1 h at 4°C. The mixture was added to pre-chilled Vero or Vero+ACE2+TMPRSS2 cells at an MOI of 0.005 and incubated at 4°C for 1 h. Cells were washed six times with chilled PBS before addition of lysis buffer and extraction of RNA using MagMax viral RNA isolation kit (Thermo Fisher Scientific) and a Kingfisher Flex 96-well extraction machine (Thermo Fisher Scientific). SARS-CoV-2 RNA was quantified by qRT-PCR using the N-specific primer/ probe set described below. Viral RNA levels were normalized to GAPDH, and the fold change was compared with isotype control mAb.

### Human ACE2 binding inhibition analysis

Wells of 384-well microtiter plates were coated with 1 μg/mL purified recombinant SARS-CoV-2 S2_Pecto_ protein at 4°C overnight. Plates were blocked with 2% non-fat dry milk and 2% normal goat serum in DPBS-T for 1 h. For screening assays, purified mAbs from microscale expression were diluted two-fold in blocking buffer starting from 10 μg/mL in triplicate, added to the wells (20 μL per well) and incubated for 1 h at ambient temperature. Recombinant human ACE2 with a C-terminal Flag tag peptide was added to wells at 2 μg/mL in a 5 μL per well volume (final 0.4 μg/mL concentration of human ACE2) without washing of antibody and then incubated for 40 min at ambient temperature. Plates were washed and bound human ACE2 was detected using HRP-conjugated anti-Flag antibody (Sigma-Aldrich, cat. A8592, lot SLBV3799, 1:5,000 dilution) and TMB substrate. ACE2 binding without antibody served as a control. The signal obtained for binding of the human ACE2 in the presence of each dilution of tested antibody was expressed as a percentage of the human ACE2 binding without antibody after subtracting the background signal. For dose–response assays, serial dilutions of purified mAbs were applied to the wells in triplicate, and mAb binding was detected as detailed above. IC_50_ values for inhibition by mAb of S2_Pecto_ protein binding to human ACE2 was determined after log transformation of antibody concentration using sigmoidal dose–response nonlinear regression analysis.

### Electron microscopy negative stain grid preparation, imaging and processing of S6_Pecto_–Fab complexes

To perform electron microscopy imaging, Fabs were recombinantly expressed and purified or produced by digesting recombinant chromatography-purified IgGs using resin-immobilized cysteine protease enzyme (FabALACTICA, Genovis). The digestion occurred in 100 mM sodium phosphate and 150 mM NaCl pH 7.2 (PBS) for around 16 h at ambient temperature. To remove cleaved Fc from intact IgG, the digestion mix was incubated with CaptureSelect Fc resin (Genovis) for 30 min at ambient temperature in PBS buffer. If needed, the Fab was buffer-exchanged into Tris buffer by centrifugation with a Zeba spin column (Thermo Fisher Scientific).

For screening and imaging of negatively-stained SARS-CoV-2 S6_Pecto_ protein in complex with human Fabs, the proteins were incubated at an Fab:spike monomer molar ratio of 4:3 for about 1 hour at ambient temperature, and approximately 3 μL of the sample at concentrations of about 10– 15 μg/mL was applied to a glow-discharged grid with continuous carbon film on 400 square mesh copper electron microscopy grids (Electron Microscopy Sciences). The grids were stained with 0.75% uranyl formate (Ohi et al., 2004). Images were recorded on a Gatan US4000 4k × 4k CCD camera using an FEI TF20 (TFS) transmission electron microscope operated at 200 keV and control with Serial EM. All images were taken at 50,000× magnification with a pixel size of 2.18 Å per pixel in low-dose mode at a defocus of 1.5 to 1.8 μm. The total dose for the micrographs was around 30e− per Å2. Image processing was performed using the cryoSPARC software package. Images were imported, CTF-estimated and particles were picked. The particles were extracted with a box size of 256 pixels and binned to 128 pixels. 2D class averages were performed and good classes selected for *ab initio* model and refinement without symmetry. Model docking to the EM map was done in Chimera (Pettersen et al., 2004). For SARS-CoV-2 S6_Pecto_ protein, the closed model (PDB:7JJI) was used and PDB:12E8 was used for the Fab (see also **Table S2** for details).

### Epitope mapping of antibodies by alanine-scanning mutagenesis

Epitope mapping was performed essentially as described previously (Davidson and Doranz, 2014) using a SARS-CoV-2 (strain Wuhan-Hu-1) spike protein NTD shotgun mutagenesis mutation library, made using a full-length expression construct for spike protein, where 215 residues of the NTD (between spike residues 9 and 310) were mutated individually to alanine, and alanine residues to serine. Mutations were confirmed by DNA sequencing, and clones arrayed in a 384-well plate, one mutant per well. Binding of mAbs to each mutant clone in the alanine scanning library was determined, in duplicate, by high-throughput flow cytometry. A plasmid encoding cDNA for each spike protein mutant was transfected into HEK-293T cells and allowed to express for 22 h. Cells were fixed in 4% (v/v) paraformaldehyde (Electron Microscopy Sciences), and permeabilized with 0.1% (w/v) saponin (Sigma-Aldrich) in PBS plus calcium and magnesium (PBS++) before incubation with mAbs diluted in PBS++, 10% normal goat serum (Sigma), and 0.1% saponin. MAb screening concentrations were determined using an independent immunofluorescence titration curve against cells expressing wild-type S protein to ensure that signals were within the linear range of detection. Antibodies were detected using 3.75 μg/mL of Alexa-Fluor-488-conjugated secondary antibodies (Jackson ImmunoResearch Laboratories) in 10% normal goat serum with 0.1% saponin. Cells were washed three times with PBS++/0.1% saponin followed by two washes in PBS, and mean cellular fluorescence was detected using a high-throughput Intellicyte iQue flow cytometer (Sartorius). Antibody reactivity against each mutant S protein clone was calculated relative to wild-type S protein reactivity by subtracting the signal from mock-transfected controls and normalizing to the signal from wild-type S-transfected controls. Mutations within clones were identified as critical to the mAb epitope if they did not support reactivity of the test MAb, but supported reactivity of other SARS-CoV-2 antibodies. This counter-screen strategy facilitates the exclusion of S protein mutants that are locally misfolded or have an expression defect.

### Selection of virus escape mutants using the S protein-expressing VSV

To screen for escape mutations selected in the presence of individual mAbs, we used a modification of the RTCA assay as recently described (Greaney et al., 2021). Fifty μL of cell culture medium (DMEM supplemented with 2% FBS) was added to each well of a 96-well E-plate to obtain a background reading. A suspension of 18,000 Vero E6 cells in 50 μL of cell culture medium was seeded per each well, and plates were placed on the analyzer. Measurements were taken automatically every 15 min and the sensograms were visualized using RTCA software version 2.1.0 (ACEA Biosciences Inc). VSV-SARS-CoV-2 virus (5,000 PFU per well, ∼0.3 MOI) was mixed with a saturating neutralizing concentration of COV2-2676 (5 μg/mL) and COV2-2489 (50 µg/mL) in a total volume of 100 mL and incubated for 1 h at 37°C. At 16 to 20 h after seeding the cells, the virus-antibody mixtures were added into 1 to 88 replicate wells of 96-well E-plates with cell monolayers. Wells containing only virus in the absence of antibody and wells containing only Vero E6 cells in medium were included on each plate as controls. Plates were measured continuously (every 15 min) for 72 h. The escape mutants were identified by unexpectedly high CPE in wells containing neutralizing antibody. To verify escape from antibody selection, isolated viruses were assessed in a subsequent RTCA experiment in the presence of 10 μg/mL (COV2-2676) and 100 µg/mL (COV2-2489) of mAb as used for the escape virus selection.

### Sequence analysis of the gene encoding S protein from S protein-expressing VSV escape mutants

To identify escape mutations present in S protein-expressing VSV mAb-selected escape variants, the escape viruses isolated after RTCA escape screening were propagated in 6-well culture plates with confluent Vero E6 cells in the presence of 10 μg/mL of the corresponding antibody. Viral RNA was isolated using a QiAmp Viral RNA extraction kit (QIAGEN) from aliquots of supernatant containing a suspension of the selected virus population. The S protein gene cDNA was amplified with a SuperScript IV One-Step RT-PCR kit (Thermo Fisher Scientific) using primers flanking the S gene. The amplified PCR product (4,000 bp) was purified using SPRI magnetic beads (Beckman Coulter) at a 1:1 ratio and sequenced by the Sanger sequence technique using primers giving forward and reverse reads of the NTD.

### MAb binding to the surface of SARS-CoV-2 infected cells

Vero E6 cells were inoculated with SARS-CoV-2 at an MOI of 0.01. At 48 h post-infection, cells were trypsinized and resuspended in a staining buffer composed of DPBS with 5% FBS, 5 mM EDTA, 0.05% NaN3. mAbs were diluted in the staining buffer and incubated with ∼3 x 10^4^ cells for 30 min at 4°C. Cells were washed twice and incubated with Alexa Fluor 647-conjugated secondary antibody (Invitrogen) diluted 1:1,000 in staining buffer for 30 min at 4°C. Cells were washed twice and fixed with 4% PFA prior to detection of fluorescence signal by flow cytometry (MacsQuant) and analysis using FlowJo software.

### Protection against wild-type SARS-CoV-2 in mice

Male and female heterozygous K18-hACE C57BL/6J mice were housed in groups of up to 5 mice per cage at 18 to 24°C ambient temperatures and 40 to 60% humidity. Mice were fed a 20% protein diet (PicoLab 5053, Purina) and maintained on a 12-h light–dark cycle (06:00 to 18:00). Food and water were available *ad libitum*. Mice (8 to 9 weeks old) were inoculated with 1 × 10^3^ PFU of SARS-CoV-2 via the intranasal route. Anti-SARS-CoV-2 human mAbs or isotype control mABS were administered 24 h before (prophylaxis) or 24 h or 48 h after (therapy) SARS-CoV-2 inoculation. Weights and lethality were monitored on a daily basis for up to 14 days after inoculation and a subset of mice were euthanized at 7 dpi and tissues were collected.

### Measurement of viral burden

For RT–qPCR, mouse tissues were weighed and homogenized with zirconia beads in a MagNA Lyser instrument (Roche Life Science) in 1 mL of DMEM medium supplemented with 2% heat-inactivated FBS. Tissue homogenates were clarified by centrifugation at 10,000 rpm for 5 min and stored at −80°C. RNA was extracted using a MagMax mirVana Total RNA isolation kit (Thermo Fisher Scientific) and a Kingfisher Flex 96-well extraction machine (Thermo Fisher Scientific). RNA was reverse transcribed and amplified using the TaqMan RNA-to-CT 1-Step Kit (ThermoFisher). RNA levels were measured by one-step quantitative reverse transcriptase PCR (qRT-PCR) assay as previously described (Hassan *et al.,* 2020). A TaqMan assay was designed to target the N gene, as previously described (Case; PMID 32838945). Specific primers and probe were used: Forward primer: ATGCTGCAATCGTGCTACAA; Reverse primer: GACTGCCGCCTCTGCTC; Probe: /56-FAM/TCAAGGAAC/ZEN/AACATTGCCAA/3IABkFQ/. N gene copy numbers were determined down to 10 copies per reaction.

### ELISA to detect recombinant human mAbs

Goat anti-human kappa cross-absorbed against mouse IgG (Southern Biotech catalog # 2061-01) and goat anti-human lambda cross-absorbed against mouse IgG (Southern Biotech catalog # 2071-01) were coated onto 96-well Nunc Maxisorp flat-bottomed plates at 2 µg/mL in coating buffer (0.1 M sodium carbonate, 0.1 M sodium bicarbonate, 0.02% NaN3, pH 9.6) overnight at 4°C. Coating buffers were aspirated, and wells were blocked with 2% BSA (blocking buffer) (Fisher Bioreagents catalog # BP1600-100), for 1 h at 37°C. Heat-inactivated serum samples were diluted in blocking buffer in a separate polypropylene plate. The plates then were washed 4× with 1× PBS + 0.05% Tween-20 (PBST) (Fisher Bioreagents catalog # BP337-100), followed by addition of 50 µL of respective serum dilutions and was incubated for 1 h at 4°C. The ELISA plates were again washed 4× in PBST, followed by addition of 50 µL of 1:2,000 Goat Anti-Human IgG Fc, Multi-Species (Southern Biotech catalog # 2014-05). Plates were incubated for 1 h at 4°C. Plates were washed with 4× PBST, followed by 100 µL of TMB-ELISA substrate (Thermo Fisher Scientific catalog # 34028) and incubated at room temperature for 3 to 5 min. Color development was observed and reactions were stopped with 50 µL of 2N sulfuric acid. Optical density (450 nm) measurements were determined using a microplate reader (Bio-Rad).

### Cytokine and chemokine protein measurements

Lung homogenates were incubated with Triton-X-100 (1% final concentration) for 1 h at room temperature to inactivate SARS-CoV-2. The samples were then analyzed by Eve Technologies Corporation (Calgary, AB, Canada). Cytokine and chemokine protein expression were determined using the Mouse Cytokine Array / Chemokine Array 31-Plex (MD31) platform. Fold-change was calculated by comparing anti-SARS-CoV-2-specific or isotype-control mAb-treated mice to naive control mice.

### Lung histology

Mice were euthanized and tissues were harvested before lung was inflation and fixation. The left lung was first tied off at the left main bronchus and collected for viral RNA analysis. The right lung was inflated with approximately with 1.2 mL of 10% neutral buffered formalin using a 3-mL syringe and catheter inserted into the trachea. The inflated lung was then kept in 40 mL neutral buffered formalin for 7 days. Tissues were embedded in paraffin, and sections were stained with hematoxylin and eosin. Tissue sections were then scanned using Hamamatsu NanoZoomer slide scanning system. Scanned image was then viewed by using the NDP view software (ver.1.2.46).

### QUANTIFICATION AND STATISTICAL ANALYSIS

Mean ± S.E.M. or mean ± S.D. were determined for continuous variables as noted. Technical and biological replicates are described in the figure legends. For analysis of mouse studies, the comparison of weight-change curves was performed using a one-way ANOVA with Dunnett’s post hoc test using Prism v.9.0 (GraphPad). Viral burden and gene-expression measurements were compared to each other or the isotype control using a one-way ANOVA with Tukey’s or Dunnett’s post hoc test, respectively, using Prism v.9.0 (GraphPad). Survival curves were estimated using the Kaplan-Meier method and differences assessed using the log-rank Mantel-Cox test and a Bonferroni correction for multiple comparisons using Prism v.9.0 (GraphPad).

---

**Figure S1.**
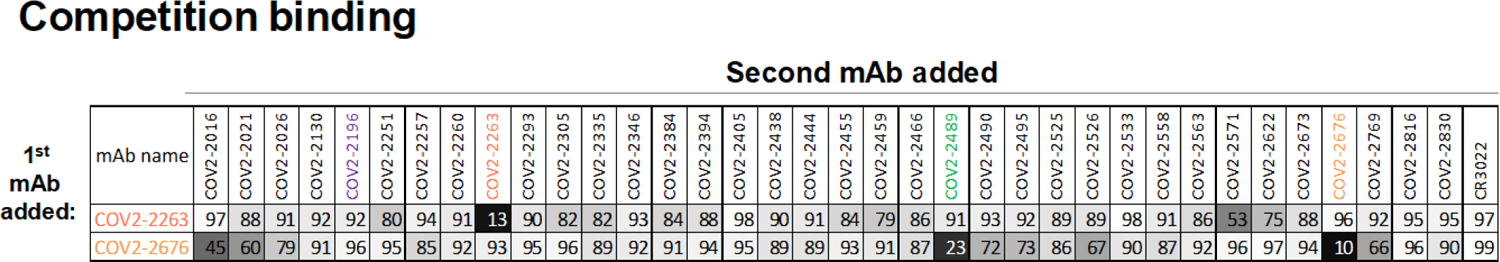
Competition binding of NTD-reactive MAbs. Competition of the panel of neutralizing mAbs with reference mAbs COV2-2676 and COV2-2263. Binding of reference mAbs to trimeric S-6P_ecto_ was measured in the presence of a saturating concentration of competitor mAb in a competition ELISA and normalized to binding in the presence of rDENV-2D22. Black indicates full competition (<25% binding of reference antibody); grey indicates partial competition (25 to 60% binding of reference antibody); white indicates no competition (>60% binding of reference antibody).

**Figure S2.**
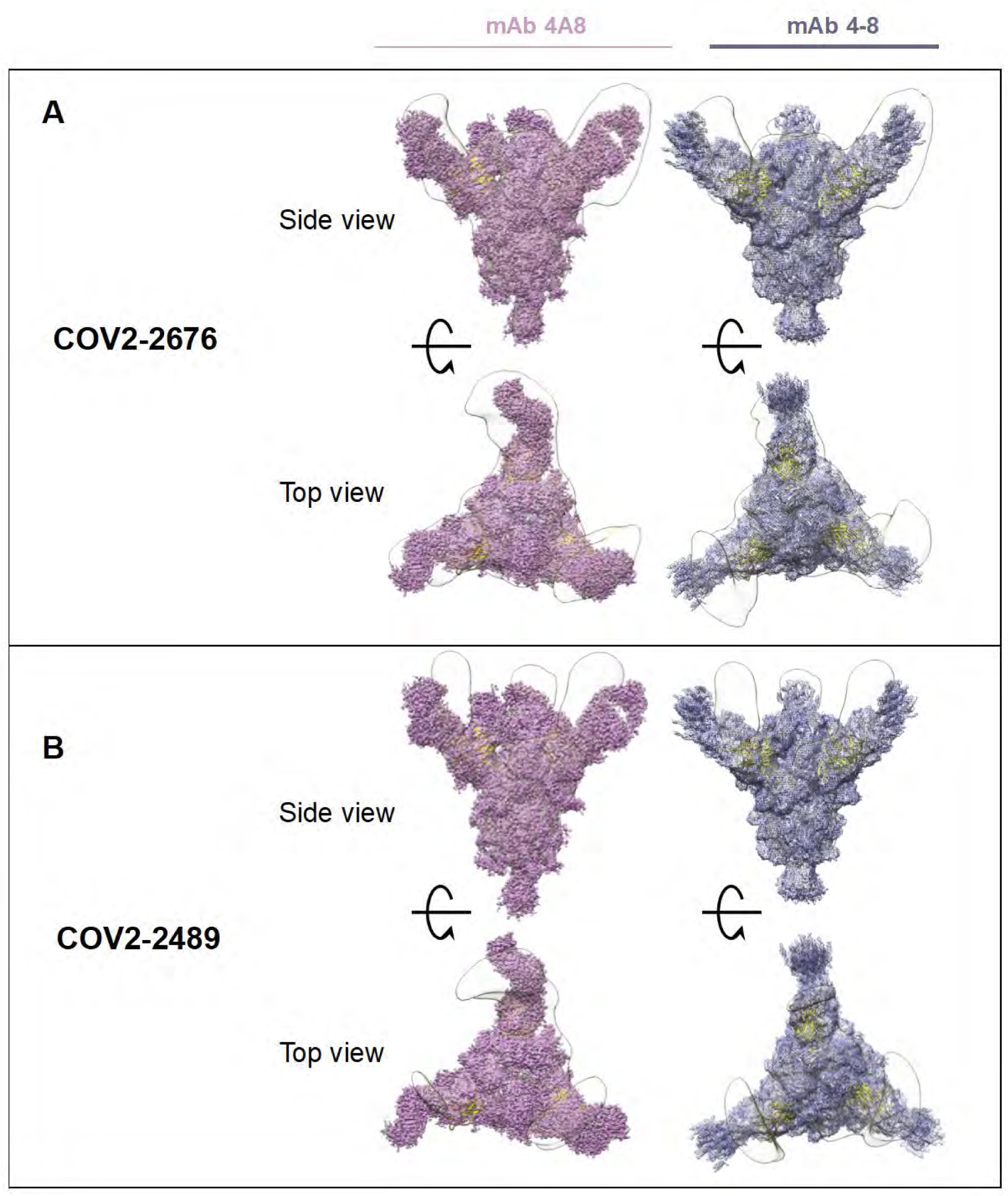
Superimposed Fab-Spike negative stain EM. **A)** COV2-2676 with mAb on the top is side view 4A8 (left), mAb 4-8 (right) and bottom is top view of the same. **B)** COV2-2489 with mAb on the top is side view 4A8 (left), mAb 4-8 (right) and bottom is top view of the same.

**Figure S3.**
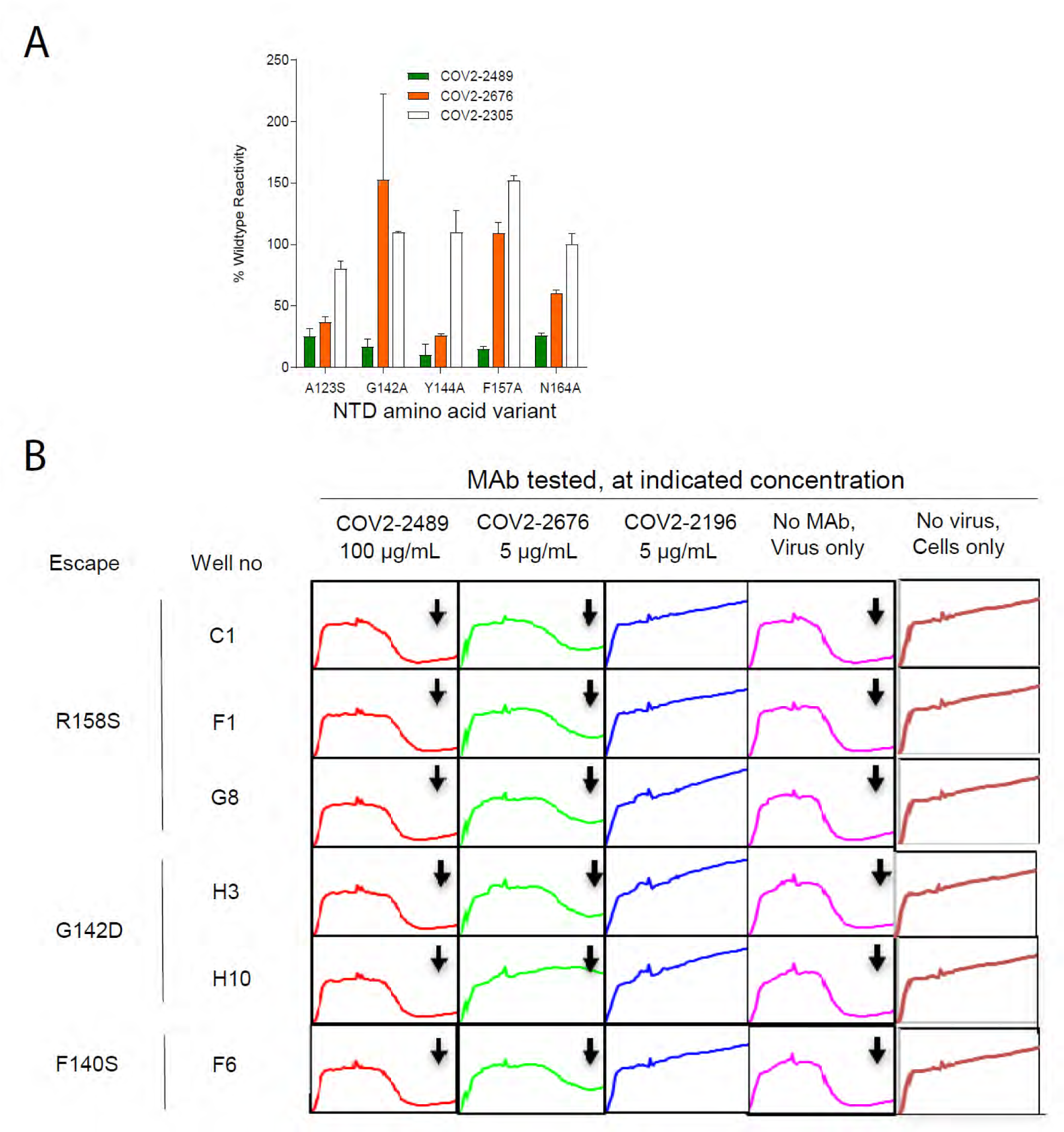
A) Identification of critical contact residues by alanine mutagenesis. Binding values for mAbs COV2-2489, −2676, and −2305. The binding values are shown as a percentage of mAb binding to wild-type (WT) SARS-CoV-2 spike protein and are plotted with the range (highest-minus lowest binding value) of at least two measurements. **B) Real-time cell analysis (RTCA) to select for spike-protein-expressing VSV viruses that escape antibody neutralization, related to** Figure 2 **B & C.** Example sensograms from individual wells of 96-well E-plate analysis showing viruses that escaped neutralization (noted with arrow) by indicated antibodies. Cytopathic effect (CPE) was monitored kinetically in Vero E6 cells inoculated with virus in the presence of a saturating concentration of antibody COV2-2489 at 100 μg/mL (red), COV2-2676 at 5 μg/mL (green) or lack of escape using RBD-specific mAb COV2-2196-blue at 5 μg/mL (blue) are shown. Uninfected cells (brown) or cells inoculated with virus without antibody (magenta) serve as controls. All curves represented show a mean of technical duplicates.

**Figure S4.**
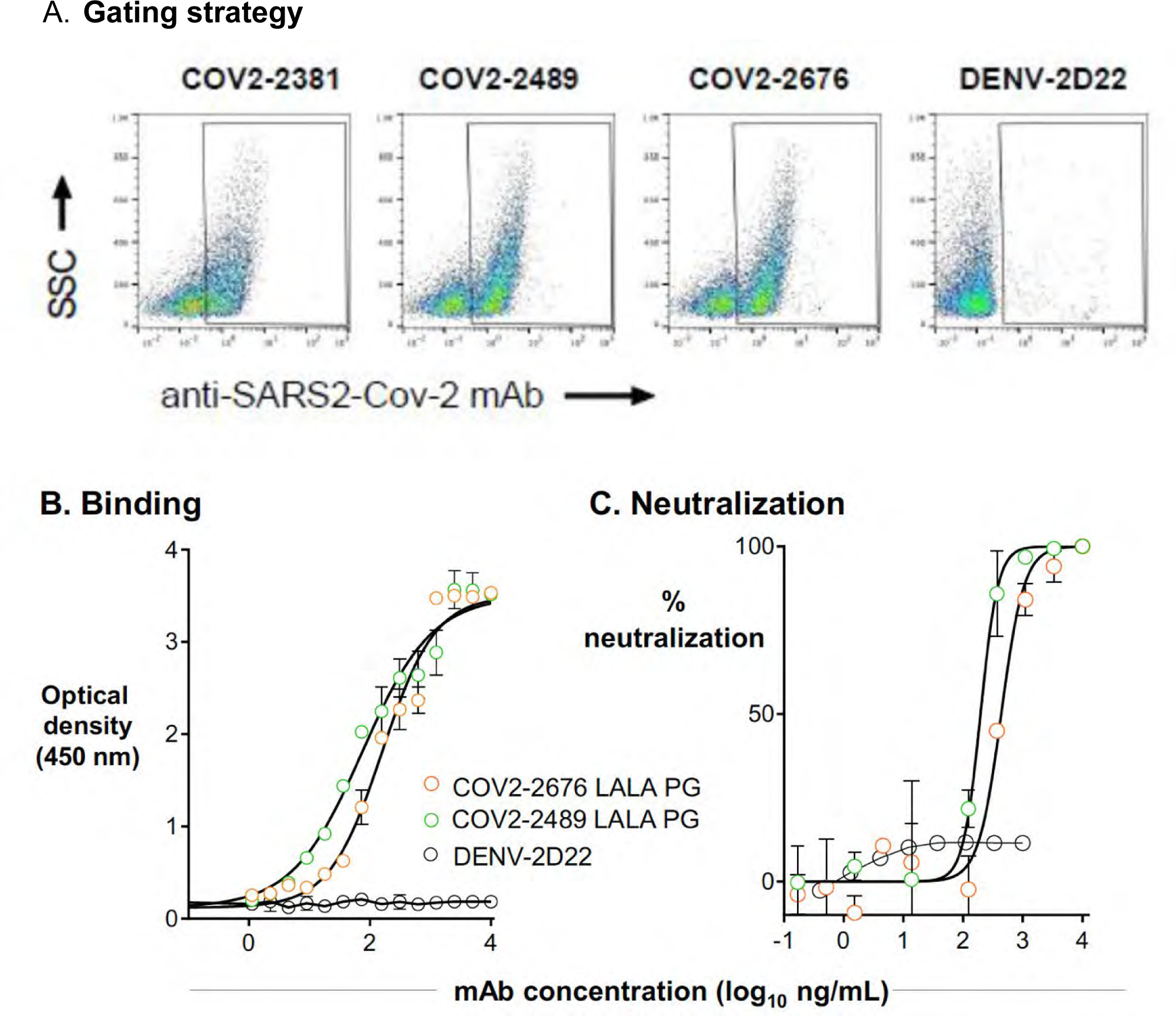
Gating strategy of Vero-E6 cells infected with SARS-CoV-2 for cell surface-displayed spike protein binding assay and ELISA and FRNT for COV2-2676 and COV2-2489 mAbs made with LALA-PG Fc variants. **A.** A representative gating strategy illustrating stained with primary COV2-2381, COV2-2676, COV2-2489 or DENV-2D22 MAb. **B.** ELISA binding of COV2-2676-LALA-PG, COV2-2489-LALA-PG or DENV-2D22 to trimeric S-6_Pecto_. Data are mean ± S.D. of technical triplicates from a representative experiment repeated twice. **C.** Neutralization curves for COV2-2676-LALA-PG, 2489-LALA-PG or DENV-2D22 using wild-type authentic SARS-CoV-2 in a FRNT assay. Error bars indicate S.D.; data represent at least two independent experiments performed in technical duplicate.

**Figure. S5A-C.**
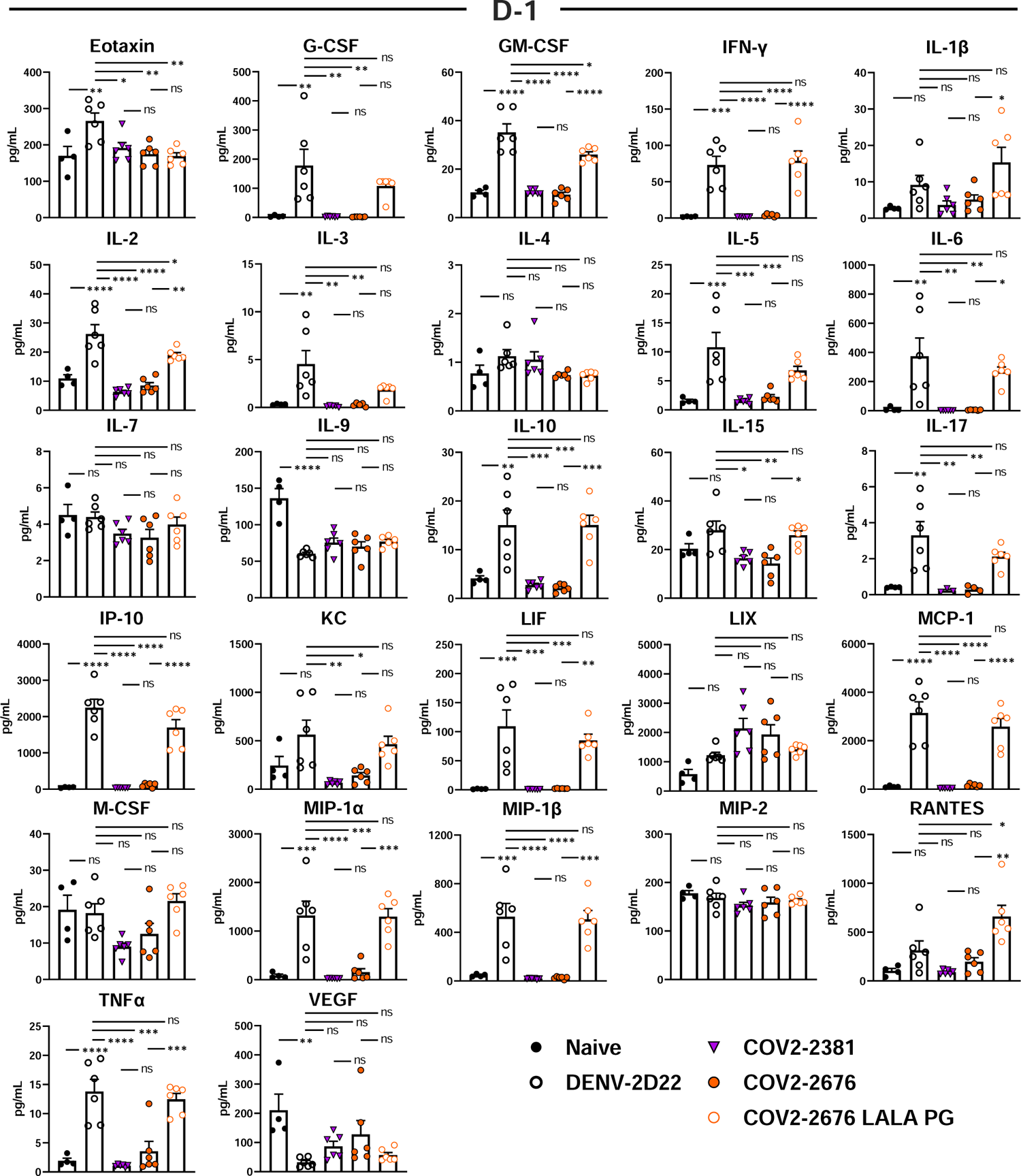
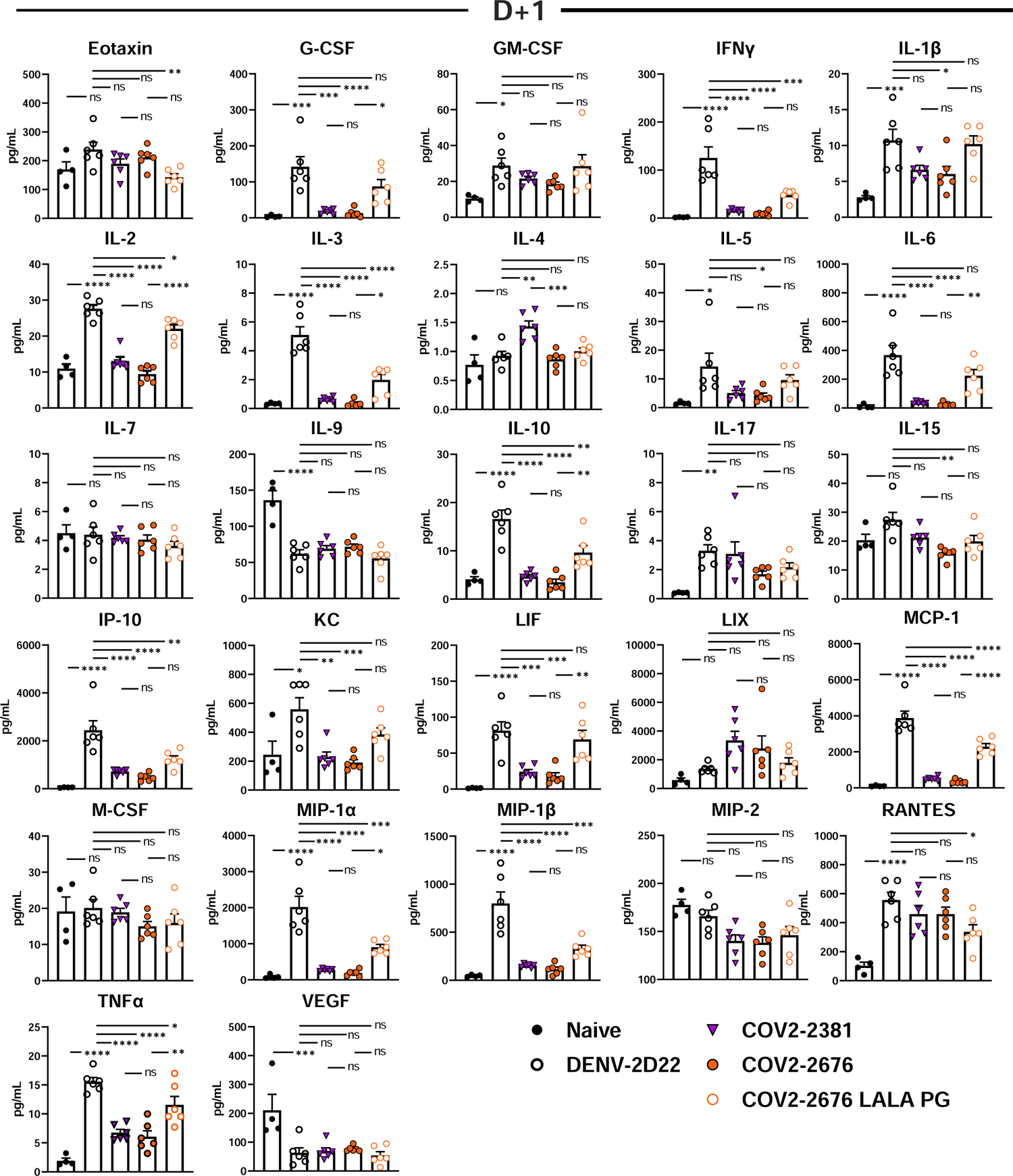
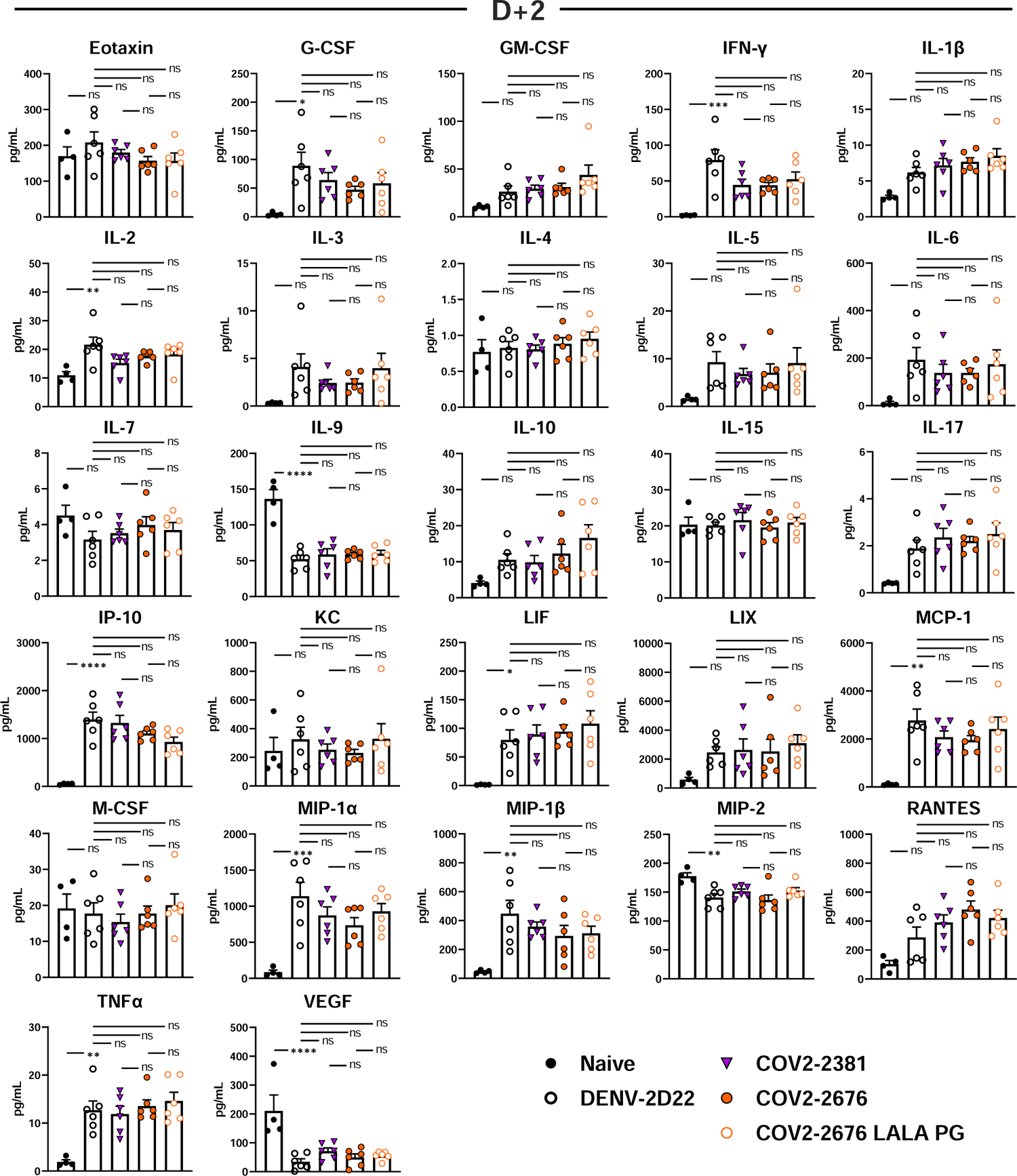
Cytokine and chemokine levels in the lungs of SARS-CoV-2 infected mice. A. Cytokine and chemokine levels in the lungs of SARS-CoV-2 infected mice at 7 dpi following d-1 treatment with isotype, COV2-2381, COV2-2676, and COV2-2676 LALA PG as measured by a multiplex platform (two independent experiments, n = 6 per group. One-way ANOVA with Tukey’s post hoc test: ns, not significant, ^∗^p < 0.05, ^∗∗^p < 0.01, ^∗∗∗^p < 0.001, ^∗∗∗∗^p < 0.0001.) B. Cytokine and chemokine levels in the lungs of SARS-CoV-2 infected mice at 7 dpi following d+1 treatment with isotype, COV2-2381, COV2-2676, and COV2-2676 LALA PG as measured by a multiplex platform (two independent experiments, n = 6 per group. One-way ANOVA with Tukey’s post hoc test: ns, not significant, ^∗^p < 0.05, ^∗∗^p < 0.01, ^∗∗∗^p < 0.001, ^∗∗∗∗^p < 0.0001.) C. Cytokine and chemokine levels in the lungs of SARS-CoV-2 infected mice at 7 dpi following d+1 treatment with isotype, COV2-2381, COV2-2676, and COV2-2676 LALA PG as measured by a multiplex platform (two independent experiments, n = 6 per group. One-way ANOVA with Tukey’s post hoc test: ns, not significant, ^∗^p < 0.05, ^∗∗^p < 0.01, ^∗∗∗^p < 0.001, ^∗∗∗∗^p < 0.0001.)

